# KAI2 regulates root and root hair development by modulating auxin distribution

**DOI:** 10.1101/539734

**Authors:** José Antonio Villaecija-Aguilar, Maxime Hamon-Josse, Samy Carbonnel, Annika Kretschmar, Karin Ljung, Tom Bennett, Caroline Gutjahr

## Abstract

Strigolactones (SLs) are endogenous signalling molecules that play important roles in controlling plant development. SL perception is closely related to that of karrikins, smoke-derived compounds presumed to mimic endogenous signalling molecules (KLs). SLs have been suggested to regulate root development. However, perception of both molecules requires the F-box protein MAX2 and the use of *max2* mutants has hampered defining the exact role of SLs in roots. Here we dissect the role of SL and KL signalling in *Arabidopsis* root development using mutants defective in the α/β hydrolase receptors D14 and KAI2, which specifically perceive SLs and KLs, respectively. Both pathways together regulate lateral root density (LRD), but contrary to previous reports, KL signalling alone controls root hair density, root hair length and additionally root skewing, straightness and diameter. Members of the SMXL protein family are downstream targets of SL (SMXL6, 7, 8) and KL (SMAX1, SMXL2) signalling. We identified distinct and overlapping roles of these proteins in the regulation of root development. Both SMAX1/SMXL2 and SMXL6/SMXL7/SMXL8 regulate LRD, confirming that SL and KL signalling act together to regulate this trait, while the KL-signalling specific SMAX1 and SMXL2 regulate all other investigated root traits. Finally, we show that KL signalling regulates root hair development by modulating auxin distribution within the root.

## INTRODUCTION

Plant roots continually integrate environmental information to make decisions about their development, allowing them to optimize their growth for optimal nutrient uptake and anchorage. This may require localized developmental responses to local environmental stimuli such as obstacles in the soil, nutrient-rich soil pockets or whole root system responses to systemic signals arriving from the shoot (Rellán-Álvarez et al., 2016). A variety of root morphological alterations are the result of different necessities of the plant. For example, nutrient deficiencies increase lateral root and root hair formation, leading to an increased root surface area and allowing plants to explore portions of soil rich in nutrients (Zhang and Forde, 2000; Ma et al., 2001; Gruber et al., 2013). The different parts of the root system may exhibit divergent sensitivities to the same factor, which indicates an autonomous control of different root system architecture traits (Jain et al., 2007; Julkowska et al., 2014). Root development of Arabidopsis is typically analyzed in seedlings germinated in a petri dish on an agar surface. In these conditions, a range of traits, such as lateral root density, primary root length, gravitropism or root hair development are easily observed. In addition, assymetric touch stimuli resulting from growth on the agar surface, cause growth behaviours such as root skewing and waving (Okada and Shimura, 1990; Vaughn and Masson, 2011), and act as an additional readout to study molecular mechanisms controlling root development.

The transmission and integration of local information within the root system and systemic information stemming from the shoot or distantly located root system portions are critical for the regulation of root system development. A number of low-molecular-weight signalling molecules (‘phytohormones’) are used to achieve these localized developmental responses as well as transmission and integration of information across long distances. The exact developmental response depends on the tissue in question, and may vary extensively in nature, both qualitatively and quantitatively, between root tissues. The phytohormone auxin plays a key role in the control of root development (Overvoorde et al., 2010). Auxin biosynthesis, transport, homeostasis, perception and signalling regulate all aspects of root formation (Muday and Haworth, 1994; Casimiro et al., 2001; Ljung et al., 2001; Friml et al., 2003; Dharmasiri et al., 2005; Cheng et al., 2007; Stepanova et al., 2008; Quint et al., 2009; Tromas et al., 2009; Schlereth et al., 2010). In addition, other plant hormones influence the role of auxin in plant development (Ross et al., 2001; Schaller et al., 2015) and more specifically in root development (Benková and Hejátko, 2009; Fukaki and Tasaka, 2009; Overvoorde et al., 2010). Strigolactones (SLs) are excellent examples for phytohormones with a modulatory role. Previous studies have suggested crosstalk between SL signalling and auxin signalling and transport in the shoot, where SLs suppress branching (Shinohara et al., 2013; Soundappan et al., 2015). SL biosynthesis and signalling mutants show an over-accumulation of the auxin efflux carrier PIN-FORMED1 (PIN1) at the plasma membrane of xylem parenchyma cells and an increase of auxin transport in the stem, which is thought to result in increased shoot branching due decreased competition for auxin among shoot meristems (Crawford et al., 2010; Shinohara et al., 2013).

SLs are thought to be predominantly synthesized in the roots (Matusova et al., 2005; Xie et al., 2010) and then transported to the shoot system (Kohlen et al., 2011). The biosynthesis pathway of SLs has been identified in several plant species (Zhang and Forde, 2000; López-Ráez et al., 2008; Seto et al., 2014). The universal SL precursor carlactone is synthesized from β-carotene by a core pathway of three enzymes; the isomerase DWARF27, and the carotenoid cleavage dioxygenases CCD7 and CCD8 (MAX3 and MAX4 in Arabidopsis; RMS5 and RMS1 in pea; D10 and D17 in rice) (Jia et al., 2017). Carlactone is then modified by a variety of enzymes, including the cytochrome P450s of the MAX1 sub-family, to create a range of active SL molecules (Yoneyama et al., 2018). SLs are then perceived and hydrolysed by the α/β hydrolase DWARF14 (D14) (Hamiaux et al., 2012; de Saint Germain et al., 2016; Yao et al., 2016; Seto et al., 2019). D14 together with the SCF^MAX2^ E3 ubiquitin ligase complex is essential to trigger SL signal transduction. Besides well established roles in suppression of shoot branching, promotion of leaf lamina extension and acceleration of leaf senescence (Booker et al., 2005; Snowden et al., 2005; Gomez-Roldan et al., 2008; Umehara et al., 2008; Yamada et al., 2014) SLs have also been suggested as important regulators of root and root hair development (Koltai et al., 2010; Kapulnik et al., 2011b; Ruyter-Spira et al., 2011; Jiang et al., 2015). However, the exact role of SLs in root development remains uncertain, due to interpretational difficulties inherent in the materials used by those studies (Waters et al., 2017). These difficulties arise from the existence of a second, closely related signalling pathway that also acts through the SCF^MAX2^ complex (Nelson et al., 2011; Waters et al., 2012b; Soundappan et al., 2015).

In this second pathway MAX2 is thought to interact with KAI2 (KARRIKIN-INSENSITIVE2), a receptor protein of the α/β hydrolase family, which is encoded by an evolutionary older paralog of D14 (Delaux et al., 2012; Toh et al., 2014; Bythell-Douglas et al., 2017). KAI2 was first identified as a receptor for karrikins, a family of butenolide compounds found in the smoke of burnt plant material (Waters et al., 2012b; Guo et al., 2013). In fire-following species, karrikins are used as a germination cue, which indicates the removal of competing plants. However, karrikins promote germination across flowering plant species most of which are not fire-following, and are thus thought to mimic the action of an endogenous plant signalling molecule, which is currently denoted KAI2-ligand (KL). Indeed, a substantial body of evidence now supports the existence of KL (Waters et al., 2015; Conn and Nelson, 2016; Sun et al., 2016). Arabidopsis *kai2* mutants in the model plant Arabidopsis have germination phenotypes but also show defects in aerial organ development such as elongated hypocotyls, aberrant coleoptile angle or leaf shape (Waters et al., 2012b; Soundappan et al., 2015; Bennett et al., 2016a)). Furthermore, in rice, *kai2*/*d14l* mutants are required for colonization by arbuscular mycorrhiza fungi (Gutjahr et al., 2015) and it has been very recently shown that KAI2 also suppresses skewing and waving of Arabidopsis ecotype Landsberg *erecta* (L*er*) growing on a tilted agar surface (Swarbreck et al., 2019).

Since KAI2 and D14 act through the same F-box protein MAX2, *max2* mutants are insensitive to both SLs and karrikins, and display the combined phenotypes of *d14* and *kai2* mutants (Nelson et al., 2011; Waters et al., 2012b; Soundappan et al., 2015; Bennett et al., 2016a). Both the D14-SCF^MAX2^ and the KAI2-SCF^MAX^ complex are thought to target related members of the SMXL (SMAX1-LIKE) family of proteins, which have weak homology to ClpB type chaperonins, for proteolytic degradation. In Arabidopsis, the targets of KL signalling are SMAX1 (SUPPRESSOR OF MAX2 1) and SMXL2 while the targets of SL signalling are SMXL6, SMXL7 and SMXL8 (SMXL6,7,8) (Stanga et al., 2013; Soundappan et al., 2015; Stanga et al., 2016). The exact molecular function of the SMXL proteins has not been defined. But SMXL6,7,8 and the protein product of their rice ortholog D53 have been associated with transcriptional regulation of a small number of target genes (Soundappan et al., 2015; Song et al., 2017). Rice D53 interacts with IPA1, a SQUAMOSA PROMOTER-BINDING FAMILY LIKE transcription factor in the regulation of shoot branching (Song et al., 2017) and stabilizes the interaction of the co-repressor TOPLESS with nucleosomes *in vitro* (Ma et al., 2001; Soundappan et al., 2015; Wang et al., 2015) suggesting that SMXLs may be directly involved in transcriptional repression. However, they have also been found to enhance PIN1 accumulation at the basal membrane of stem xylem parenchyma cells and auxin transport (Liang et al., 2016).

The close relationship between SL and KL signalling also extends to the biochemical structure of the ligands. The stereochemistry of SLs is complex, with most SLs having two chiral centres, such that SLs can theoretically exist in four stereo-conformations. Only two of these are naturally occurring (the so-called *5DS* and *4DO* conformations), and there is genetic evidence that D14 can bind and respond to either type (Scaffidi et al., 2014). It is assumed that KAI2 does not bind to naturally occurring SLs, but it does bind and hydrolyse the non-natural *ent-5DS* stereoisomers of SLs (Waters et al., 2015). This suggests KL is a molecule of similar size and structure to SLs, but mutations in SL synthesis enzymes do not cause *kai2*-like phenotypes, so KL is probably not derived from the SL biosynthesis pathway (Bennett et al., 2006; Scaffidi et al., 2013). Confoundingly, the major commercially available strigolactone analogue *rac*-GR24 contains a racemic mixture of SL stereoisomers, and therefore activates the signalling of both SL and KL pathways, creating a legacy of interpretational problems in past studies (Scaffidi et al., 2014; Waters et al., 2017). These problems have been exacerbated by the prolific use of the *max2* mutants, which are defective in both pathways, across many studies. This may have historical reasons as *max2* Arabidopsis mutants were available prior to *d14* and *kai2* mutants. However, if only *max2* mutants are used, these phenotypes cannot be reliably attributed to either SL or KL signalling. The interpretational problems caused by the use of *rac*-GR24 and *max2* mutants have led to confusion about the role of SLs in, for instance, seedling morphogenesis (Tsuchiya et al., 2010) and drought stress (Bu et al., 2014; Van Ha et al., 2014). As discussed above, these same problems have caused a lack of clarity about the role of SLs in root development (Waters et al., 2017).

In this study, we aimed at delineating the roles of SLs and KLs in controlling root development in Arabidopsis. Using a wide range of mutants, we comprehensively re-assessed the phenotypes caused by defective SL or KL signalling. We demonstrate that several aspects of root development previously proposed as regulated by SL signalling are instead controlled by KL signalling. We show that both SL and KL signalling regulate lateral root density, while KL signalling alone controls root skewing, root straightness and diameter. In addition, KL signalling emerges as a key regulator of root hair development. We further dissect the downstream signalling mechanisms of SLs and KL signalling using mutant analyses. We show that contrary to previous suggestions (Swarbreck et al., 2019), both SL and KL signalling likely act in a canonical manner in the root system, with SMAX1, SMXL2 being targets of KL perception and SMXL6, SMXL7, SMXL8 being targets of SL perception. Finally, we present evidence that KAI2 signalling regulates root development by modulating auxin transport and distribution within the root system, echoing the role of SLs in regulating auxin distribution within the shoot.

## RESULTS

### SL signalling has minor effects on seedling root architecture

Primary root length, lateral root density and root hair length have previously been observed to respond to *rac*-GR24 treatment and to mutation of *MAX2* (Kapulnik et al., 2011b; Kapulnik et al., 2011a; Ruyter-Spira et al., 2011; Jiang et al., 2015), which both affect SL as well as KAR signalling. To re-assess the specific role of SL in root development, we re-examined these phenotypes in mutants specifically affected in SL biosynthesis or perception, namely the SL biosynthesis mutants *max3-9, max4-5* and *max1-1* (here arranged in pathway order) and the SL receptor mutant *d14-1*.

Primary root length (PRL) and lateral root density (LRD) were determined in Leeds [L] as well as Cambridge [C]. In [L], PRL was reduced in all SL biosynthesis mutants across 5 independent experiments, though *d14-1* behaved more variable, but averaged across experiments it also had reduced PRL relative to Col-0 (Figure 1A, Supplementary Figure 1A). In [C], we examined the phenotype of *d14-1* and *max4-5* in 5 independent experiments and found little evidence for differences in PRL (Supplementary Figure 1B). Conversely, we observed only small increases in LRD for *max3-9, max4-5, max1-1* and *d14-1* relative to Col-0 and only in 2/5 experiments conducted in [L] (Figure 1B; Figure 2B). In 3/5 there were no statistically significant differences from wild-type, and averaged across experiments, there was no clear difference between SL mutants and wild-type (Supplementary Figure 1C). However, in [C] *max4-5* displayed a clear increase in LRD as compared to Col-0 in 5/5 experiments (Supplementary Figure 1D), but this increase was not as large as for *max2* mutants (Supplementary Figure 1D). In 3/6 experiments, *d14-1* was highly variable with a significantly lower LRD than Col-0, and significantly higher LRD in the other half of the experiments (Supplementary Figure 1D). Thus, consistent with previous reports (Ruyter-Spira et al., 2011), and data from other species (Sun et al., 2015; Marzec et al., 2016), we found that SL signalling has subtle effects on PRL and LRD in Arabidopsis, which appear to be sensitive to small differences in growth conditions, given the slightly different phenotypes in [L] or [C].

**Figure 1.**
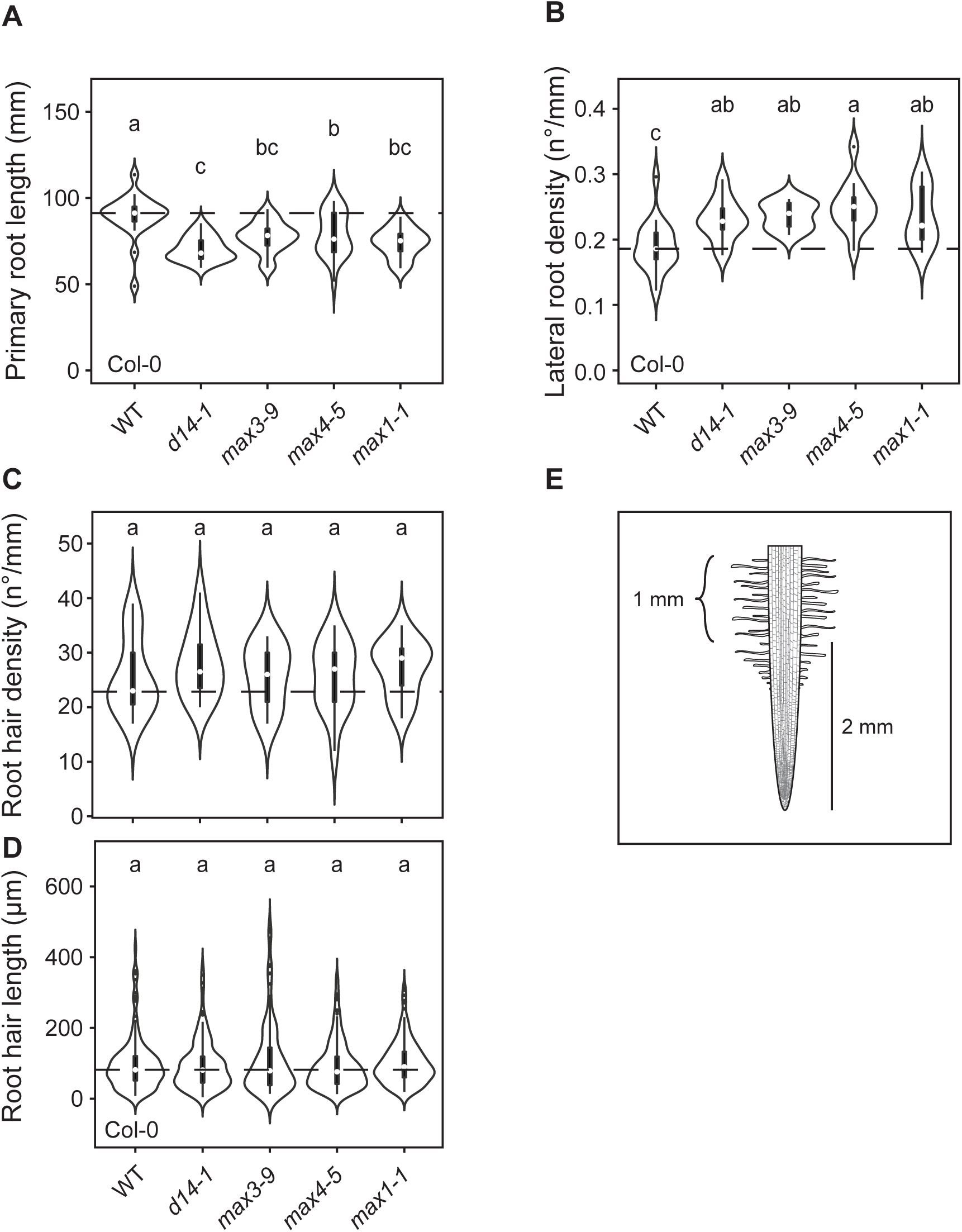
Strigolactone signalling regulates primary root length and lateral root density. **(A)** Primary root length, **(B)** lateral root density, **(C)** root hair density, and **(D)** root hair length in WT (Col-0) Arabidopsis, the strigolactone perception mutant *d14-1* and the strigolactone biosynthesis mutants *max3-9, max4-5* and *max1-1* (arranged in pathway order). **(E)** Diagram showing the primary root zone used for root hair phenotyping (curly bracket). Root hair density and length were quantified for 1 mm primary root length between 2 and 3 mm from the root tip. The outline of the violin plots represents the probability of the kernel density. Black boxes represent interquartile ranges (IQR), with the white dot representing the median; whiskers extend to the highest and lowest data point but no more than ±1.5 times the IQR from the box; outliers are plotted individually. Different letters indicate different statistical groups (ANOVA, posthoc Tukey, **(A)** F_4,111_ = 11.81; p≤ 0.001, **(B)** F_4,58_ = 5.626; p≤ 0.001, **(C)** F_4,65_ = 0.242; p≤0.05, **(D)** F_4_, _718_ = 1.291; p≤0.05).

**Figure 2.**
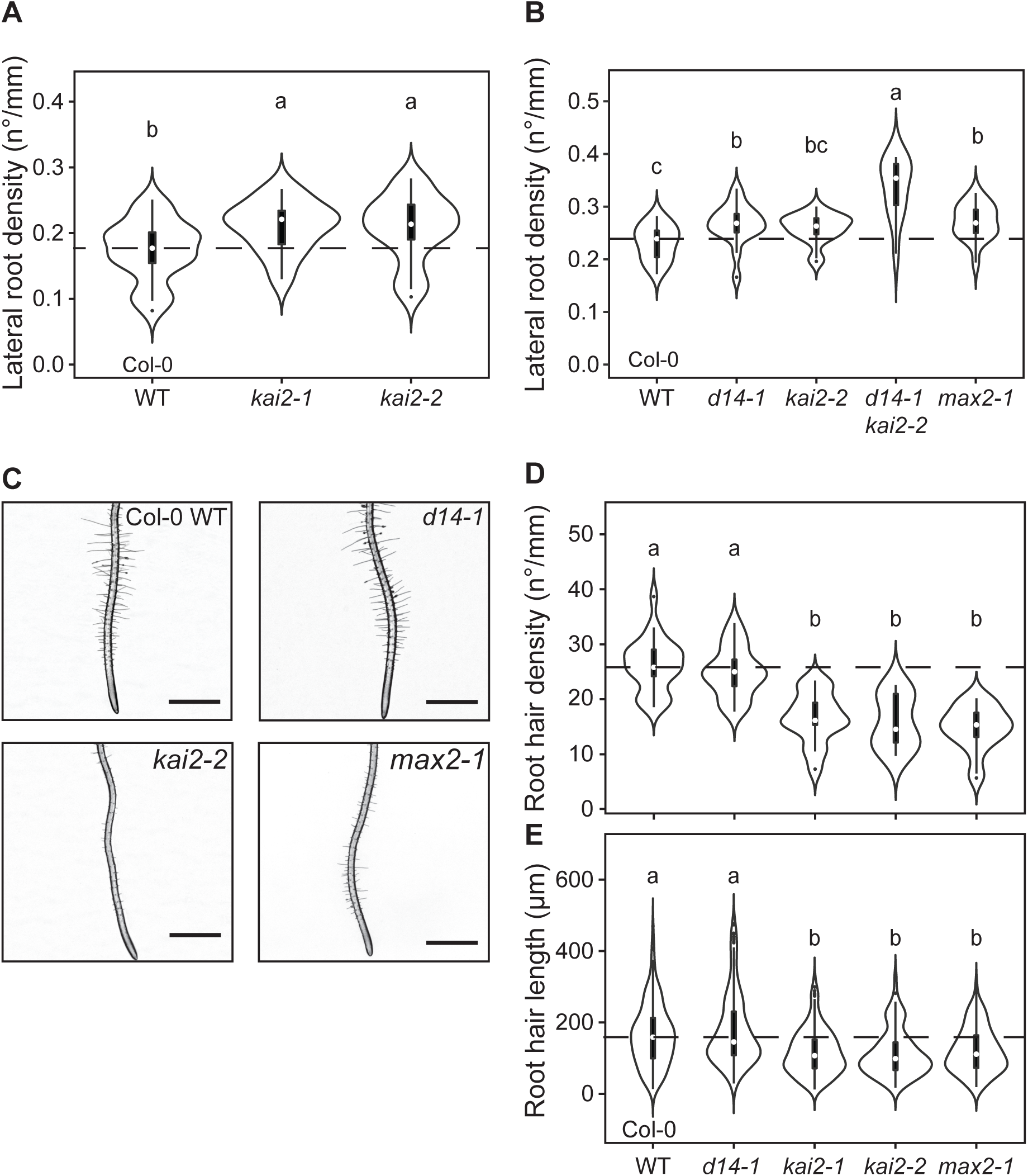
KL perception mutants are impaired in lateral root and root hair development. **(A)** Lateral root density in WT (Col-0) Arabidopsis and two *kai2* alleles. (**B-D**) Lateral root density (**B, D, E**), root hair density (**C**) and root hair length (**D**) in WT (Col-0) Arabidopsis and *d14-1 kai2-2* double mutants, with relevant single mutant controls. **(C)** Representative images of root hair phenotypes of the indicated genotypes. Scale bar, 1 mm. The outline of the violin plots represents the probability of the kernel density. Black boxes represent interquartile ranges (IQR), with the white dot representing the median; whiskers extend to the highest and lowest data point but no more than ±1.5 times the IQR from the box; outliers are plotted individually. Different letters indicate different statistical groups (ANOVA, posthoc Tukey, **(A)** F_2,79_ = 5.29; p≤ 0.01, **(B)** F_4,80_ = 15.29; p≤ 0.001, **(D)** F_4,88_ = 28.9; p≤0.001), **(E)** F_4,825_ = 23.43; p≤ 0.001).

We next examined root hair formation in the suite of SL biosynthesis and perception mutants. Contrary to previous reports (Kapulnik et al., 2011b) found that neither root hair density (RHD) nor root hair length (RHL) are statistically different between Col-0 and any of the SL synthesis or signalling mutants (Figure 1C, D). Indicating, that previously observed root hair phenotypes of *max2* mutants are caused by defects other than SL signalling, for example in KL signalling.

### KAI2 modulates lateral root density together with D14

The mutant phenotypes present in SL-specific biosynthesis and perception mutants are insufficient to account for previously described effects of *max2* on root development. Since, in the shoot, the *max2* phenotype is essentially a hybrid of *d14* and *kai2* phenotypes (Bennett et al., 2006; Soundappan et al., 2015), we reasoned that the absence of KL signalling might contribute for the observed phenotypes in *max2*. We tested this using two allelic *kai2* mutants (*kai2-1, kai2-2*) in the Col-0 background. As with SL mutants, PRL was highly variable, being larger than Col-0 in some experiments and smaller in others, but on average, there was little difference from Col-0 for any allele, across 4-7 experiments (Supplementary Figure 2A). This confirmed our observations of *max2-1*, which might have a higher or lower PRL relative to Col-0 in any given experiment, but when averaged across 5 experiments performed in [L] it was not different from Col-0 (Supplementary Figure 2A). The effects of *kai2* alleles on LRD were clearer, with increases of 10-25% relative to the wild-type and this was consistently observed across 4-7 experiments at 10 days post germination (dpg) (Figure 2A; Supplementary Figure 2B). A time-course experiment suggests that this phenotype is established early in development, as it is more clearly visible at 6 dpg than at 10 dpg, by which time Col-0 has begun to ‘catch up’. This is different for SL biosynthesis and perception mutants, which if anything lags behind Col-0 in LR emergence, and only have equal or more LRs starting from 10 dpg. (Supplementary Figure 2C). Together this suggests that SL and KL signalling both regulate LRD in Arabidopsis, but given the differences in timing of phenotype expression maybe at different developmental stages. Consistent with this hypothesis LRD phenotypes of *max2-1* were stronger as compared to *max4-5* in [C] (Supplementary Figure 1C). In [L], the *max2-1* LRD phenotype was approximately the same as for *kai2* mutants, which is consistent with the lack of expression of the SL LRD phenotype under these conditions.

We further tested the idea that both KL and SL signalling contribute to regulating LRD by examining *d14 kai2* double mutants. We observed a very strong and consistent increase in LRD in *d14-1 kai2-2* in comparison to Col-0, *d14-1* and *kai2-2*, in each of 4 independent experiments under the [L] conditions (Figure 2B). The increase in LRD was always greater in *d14-1 kai2-2* than in *max2-1* (Figure 2B). We previously observed a similar discrepancy in all shoot phenotypes examined (Bennett et al., 2016a). The point mutation in *max2-1* leads to an exchange of aspartic acid 581 to asparagine (Stirnberg et al., 2002) making it likely that *max2-1* is not a null allele. However, we cannot rule out the alternative possibility that there are MAX2-independent effects of D14 and KAI2 on root and shoot development. Nevertheless, it is now clear that the effects of *max2* mutations on PRL and LRD are attributable to the loss of both SL and KL signalling.

### KAI2 unambiguously regulates root hair development

We also examined root hair development in *kai2* mutants. Both RHD and RHL were significantly decreased for *kai2-1, kai2-2* and *max2-1* in the Col-0 background (Figure 2C-E). We observed exactly the same phenotypes for two allelic *kai2* mutants in the Ler background (Supplementary Figure 2D, E). Thus, the root hair phenotypes previously observed in *max2* mutants and attributed to lack of SL signalling are actually caused by a lack of KL signalling.

### The KAI2-MAX2 module regulates root skewing and waving

In addition to lateral root and root hair phenotypes, we observed that *kai2* mutants display increased skewing along the surface of vertically-oriented agar plates, in both Ler and Col-0 ecotypes (Figure 3, Supplementary Figure 3), consistent with a recent report (Swarbreck et al., 2019). This right-handed skewing is a well-established effect of growing Arabidopsis roots on the surface of agar plates, and probably arises from a combination of circumnutation and thigmotropic responses (Oliva and Dunand, 2007; Roy and Bassham, 2014). The effect of *kai2* on skewing is particularly pronounced in Ler, which has a stronger basal level of skew than Col-0, but is also clearly present in *kai2* mutants in the Col-0 background (Figure 3B-D; Supplementary Figure 3B). Increased skewing is also observed in *max2* mutants, but not in SL biosynthesis mutants, nor *d14* (Figure 3B, D; Supplementary Figure 3B, C, F). The increased skewing phenotype of the *d14-1 kai2-2* double mutant in the Col-0 background is equal to *kai2-2*, indicating that strigolactone perception is not involved in regulating root growth direction (Figure 3C).

**Figure 3.**
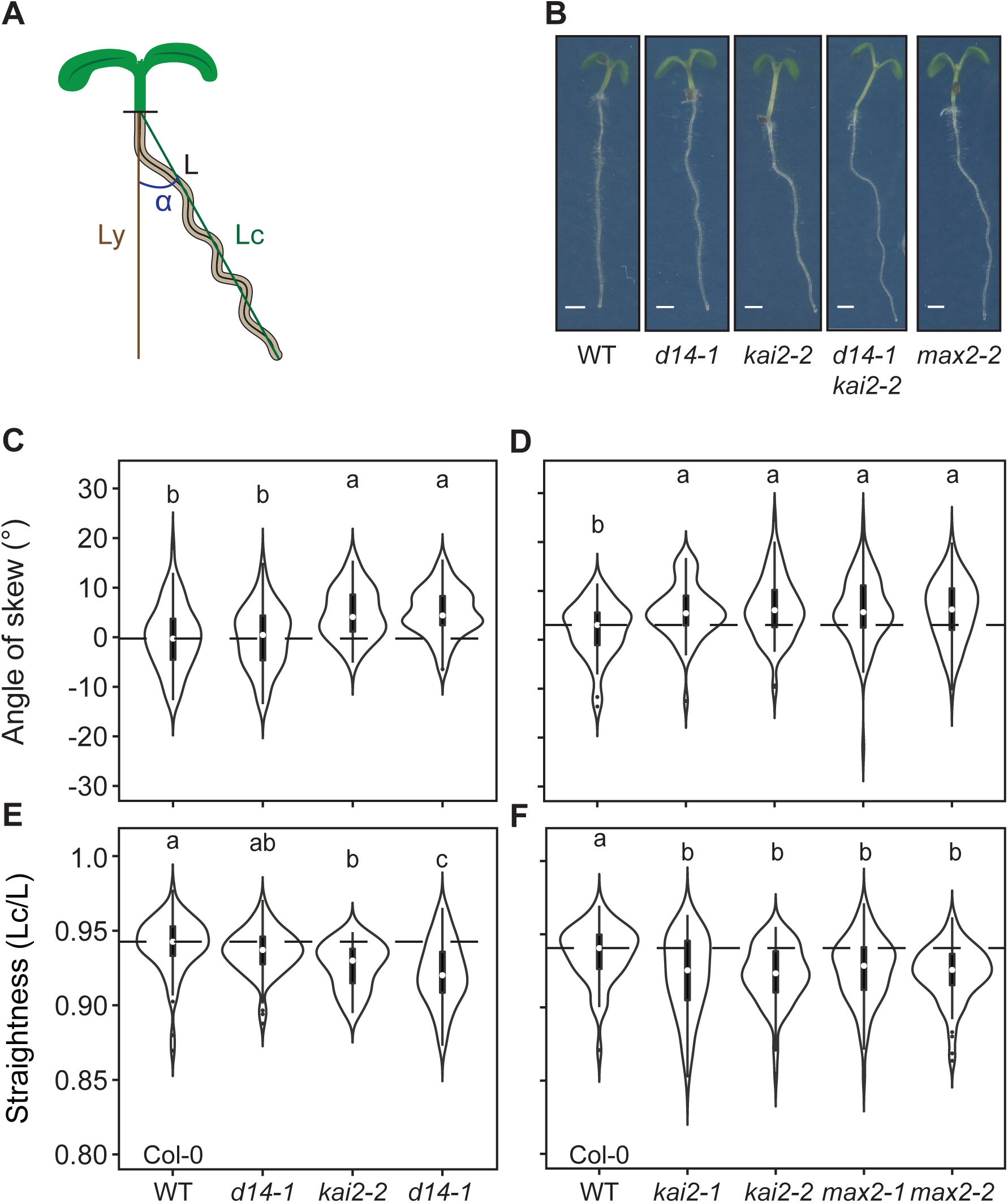
KL perception mutants display exaggerated skewing and waving. **(A)** Diagram showing how skewing-angle and root straightness were quantified. Skewing was quantified by measuring the angle between the vertical axis (Ly) defined as 0°, and the root tip. Right or left skewing is indicated by positive or negative values, respectively. Straightness was calculated as the ratio of the straight line between the hypocotyl-root junction and the root tip (green line, Lc) and the total root length (L). **(B)** Images of representative 5-days-old seedlings of the indicated genotypes. Scale bars, 1 mm. **(C**, **D)** Root skewing and **(E** and **F)** root straightness of the indicated genotypes. The outline of the violin plot represents the probability of the kernel density. Black boxes represent interquartile ranges (IQR), with the white dot representing the median; whiskers extend to the highest and lowest data point but no more than ±1.5 times the IQR from the box; outliers are plotted individually. Different letters indicate different statistical groups (ANOVA, posthoc Tukey, **(C)** F_3,315_ = 16.08; p≤ 0.001, **(D)** F_4,347_ = 4.762; p≤ 0.001, **(E)** F_3,315_ = 13.62; p≤0.001), **(F)** F_4,347_ = 4.281; p≤ 0.001).

The increased skewing phenotype in the *kai2* and *max2* mutants is accompanied by increased root waving, which can be measured as a decrease in root ‘straightness’ (Figure 3A, E, F, Supplementary Figure 3D, E, G). Again, this decreased straightness phenotype is not observed in *d14-1* or SL biosynthesis mutants (Figure 3E, Supplementary Figure 3E-G). However, the skewing and waving phenotypes seen in *kai2* mutants are to some extent genetically separable, as can be observed when plants are grown at different angles. Growth on plates inclined at 45° increases skewing specifically in Ler wild-type to the level of *kai2-2* mutants but does not trigger changes in skewing in the Col-0 wild-type or any of the mutants in both ecotypes (Supplementary Figure 3B, C). In contrast, the waving of all genotypes increases at 45°, with a concomitant decrease in straightness without considerably changing the differences observed between genotypes at 90° (Supplementary Figure 3D, E). In summary, root skewing and waving represent novel phenotypes regulated by KL but not by SL signalling in Arabidopsis roots.

### KAI2 and D14 regulate root epidermal cell elongation

Skewing is often associated with epidermal cell file rotation (Roy and Bassham, 2014). The degree of epidermal cell file rotation is quantified by counting the number of epidermal cells, which are crossed by a vertical line of a given length along the root. To determine whether skewing of KL receptor mutants is associated with cell file rotation (Wang et al., 2011), we quantified the number of epidermal cells per millimetre between millimetre 2 and 3 above the root tip. The number of cells per millimetre was indeed higher in *kai2* and *max2* mutants in the Col-0 and in the Ler ecotype (Figure 4A, Supplementary Figure 4A). However, a careful microscopic inspection of the root surface of *kai2* and *max2* mutants revealed that the epidermal cells did not show any signs of rotation; they grew in a perfect vertical orientation (Figure 4B, Supplementary Figure 4B). This is in contrast to the results of Swarbreck et al (2019), who observed increased cell file rotation in *kai2* and *max2* at a 45° growth angle. Since at a 90° growth angle we observed a skewing phenotype but no cell file rotation, we conclude that there is likely no connection between cell file rotation and skewing in *kai2* mutants. We measured the epidermal cell length and determined that epidermal cells of *kai2* and *max2* are shorter than those of the wild-type (Figure 4C, Supplementary Figure 4C). Interestingly, also *d14* displayed the short epidermal cell phenotype and this phenotype was not enhanced in the *d14 kai2* double mutant (Figure 4A, C; Supplementary Figure 4A, C). Although the exact cause of the shorter epidermal cells in KL as well as SL perception mutants is unknown, this genetic discrepancy between epidermal cell length and skewing phenotypes suggests that epidermal cell length is not directly related to skewing. Furthermore, the phenotype is non-additive suggesting that KAI2 and D14 act on an equivalent target for epidermal cell elongation.

**Figure 4.**
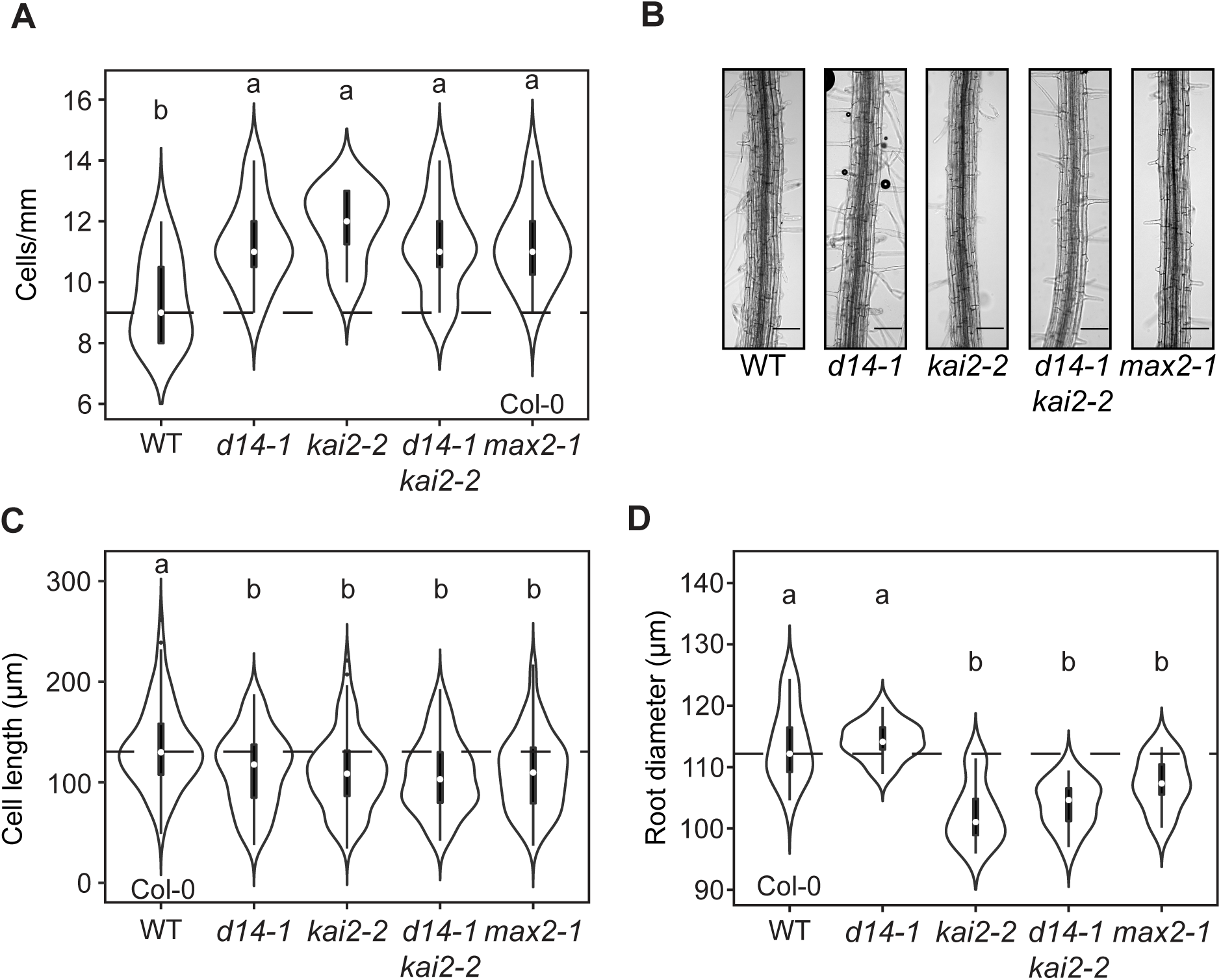
KL perception mutants exhibit decreased epidermal cell lengths and root diameter. **(A)** Number of root epidermal cells per mm of the indicated genotypes. **(B)** Images of representative roots between 2 and 3 mm from the root tip from 5-days-old seedlings of the indicated genotypes. Scale bars, 0.1 mm. **(C)** Root cell length and **(D)** root diameter of the indicated genotypes. The outline of the violin plot represents the probability of the kernel density. Black boxes represent interquartile ranges (IQR), with the white dot representing the median; whiskers extend to the highest and lowest data point but no more than ±1.5 times the IQR from the box; outliers are plotted individually. Different letters indicate different statistical groups (ANOVA, posthoc Tukey, **(A)** F_4,52_ = 4.715; p≤ 0.01, **(C)** F_3,392_ = 10.64; p≤ 0.001, **(D)** F_4,50_ = 15.95; p≤0.001).

We also noticed that *kai2* mutants in both the Col-0 and Ler background had thinner primary roots than wild-type. Quantification of root diameter at 2.5 mm above the root tip confirmed that the primary roots of *kai2* and *max2* mutants but not of the Col-0 *d14* mutant are thinner than those of the wild type (Figure 4D, Supplementary Figure 4D). This indicates that the regulation of root thickness is specific to KL signalling. It has been speculated that a smaller root cell diameter may cause tissue tensions leading to skewing (Swarbreck et al., 2019). However, we could genetically separate the thin root diameter from skewing (see below; Figure S6E), indicating that the two phenotypes are likely unrelated.

### Distinct and overlapping roles for SMXL proteins in root system and root hair development

We next assessed the role of *SMXL* genes in control of root development downstream of *KAI2* and *D14*. Previous results showed that the *max2* LRD phenotype was suppressed in a *smxl6 smxl7 smxl8* background but not in a *smax1* background (Soundappan et al., 2015), suggesting that the *max2* phenotype might arise from excess SMXL6, SMXL7 and SMXL8 (hereafter SMXL678) protein accumulation. However, care is needed in interpreting these data, since *smxl678* is also completely epistatic to *max2* in every other SL phenotype assessed (Soundappan et al., 2015; Wang et al., 2015). Thus, while it is clear that SMXL678 regulate lateral root development, the epistasis means that it is not safe to infer that SMXL678 accumulation is the sole cause of the LRD phenotype seen in *max2*. Indeed, we found that the combined loss of *SMAX1* and *SMXL2* was as efficient in suppressing the *max2* phenotype as loss of *SMXL678* (Figure 5A; Supplementary Figure 5A). Intriguingly, neither *smax1-2 smxl2-1* nor *smxl6-4 smxl7-3 smxl8-1* (hereafter *smxl678*) appeared completely epistatic to *max2* in the context of LRD. Although *smax1-2 smxl2-1* suppressed the *max2-1* phenotype to the wild-type level, *smax1-2 smxl2-1* had a significantly lower LRD than *max2-1 smax1-2 smxl2-1* in 3/4 experiments (in the fourth experiment, the genotypes had the same LRD; Supplementary Figure 5A). In a time course experiment, the phenotype of *smxl678* was less clear cut at 10dpg: in only 1/7 experiments was there a statistically significant difference between *smxl678* and *smxl678 max2-1*. However, at 6 dpg, the difference was clearer, and *smxl678* has a statistically significantly lower LRD than *smxl678 max2-1* (Supplementary Figure 5B). The most straightforward explanation for these results is that the *max2* LRD phenotype arises from the accumulation of both SMAX1/SMXL2 and SMXL678, and that neither *smax1 smxl2* nor *smxl678* alone are epistatic to the *max2* LRD phenotype as a result. However, in purely quantitative terms, both *smxl* mutant combinations can suppress the LRD phenotype of *max2-1* to the wild-type level. This confirms our observations regarding the LRD of *d14* and *kai2* mutants indicating that SL and KL signalling most likely act together in the regulation of LR development: SL signalling by promoting SMXL678 turnover, and KL signalling by promoting SMAX1 SMXL2 turnover.

**Figure 5.**
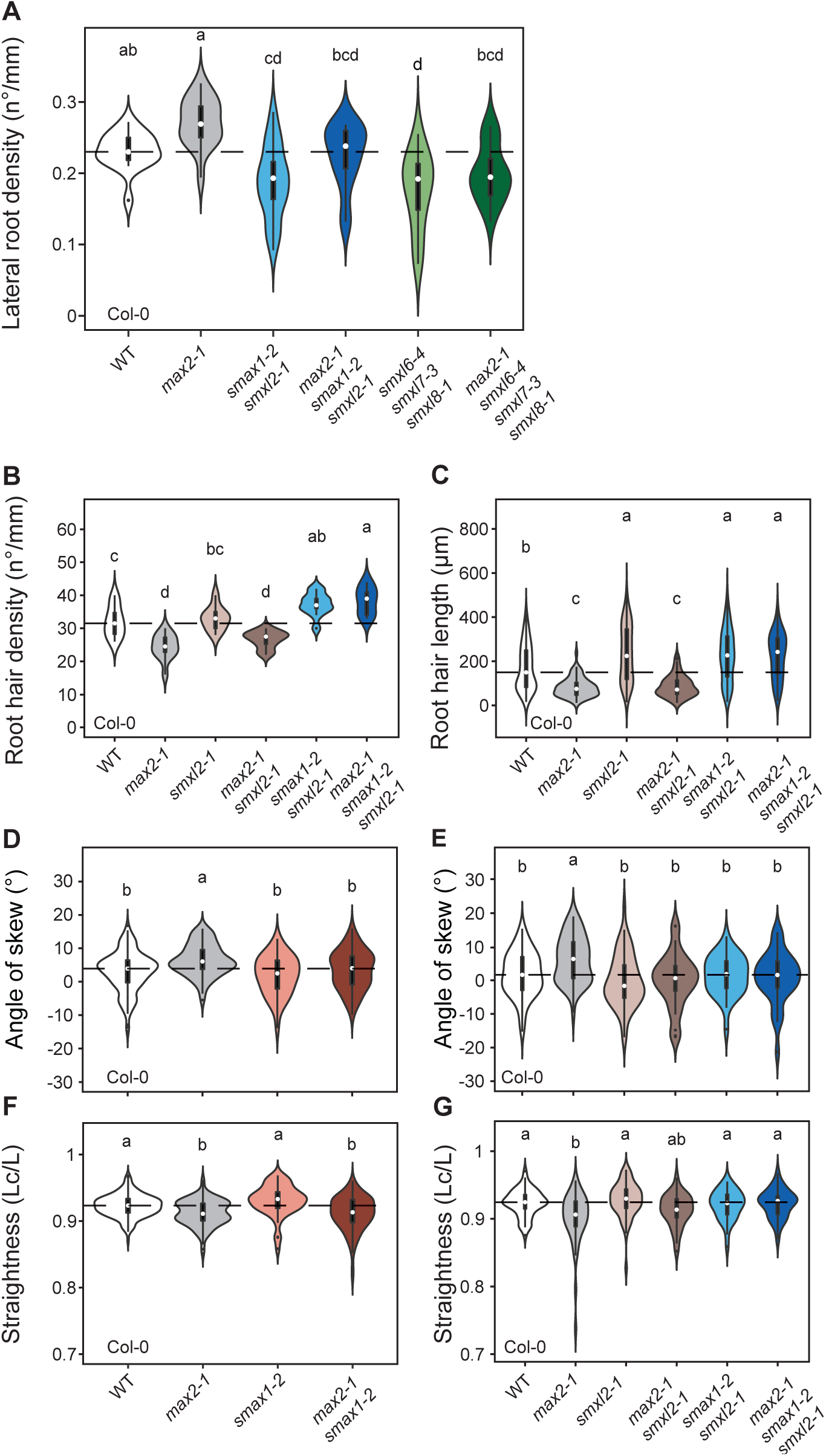
SMAX1 and SMXL2 regulate multiple aspects of root development. **(A)** Lateral root density, **(B)** root hair density, **(C)** root hair length, **(D, E)** root skewing and **(F** and **G)** root straightness of the indicated genotypes. The outline of the violin plot represents the probability of the kernel density. Black boxes represent interquartile ranges (IQR), with the white dot representing the median; whiskers extend to the highest and lowest data point but no more than ±1.5 times the IQR from the box; outliers are plotted individually. Different letters indicate different statistical groups (ANOVA, posthoc Tukey, **(A)** F_5,90_ = 10.62; p≤ 0.001, **(B)** F_5,59_ = 22.3; p≤ 0.001, **(C)** F_5,639_ = 49.95; p≤0.001, **(D)** F_3,345_ = 7.612; p≤0.001, **(E)** F_5,259_ = 5.051; p≤0.001, **(F)** F_3,440_ = 16.32; p≤0.001, **(G)** F_5,261_ = 6.57; p≤0.001).

Investigating RHD, RHL, root skewing and straightness in *smxl* mutants resulted in phenotypes consistent with the attribution of these traits to KL signalling. For RHD and RHL, the SL repressor mutants *smxl678* and *max2-1 smxl678* had the same phenotypes as Col-0 and *max2-1* respectively (Supplementary Figure 5C, D). Conversely, we found that the KL repressor mutants *smax1-2 smxl2-1* together suppressed the phenotype of *max2-1*, consistent with our observation that *kai2* and not *d14* mimicked the decreased RHD and RHL in *max2* (Figure 5B, C; Supplementary Figure 5E, F).

Interestingly, for skewing, *smax1* or *smxl2* were both independently sufficient to suppress *max2* (Figure 5D, E), indicating that skewing may be either very sensitive to the stoichiometry of SMXL proteins or that SMAX1 and SMXL2 may promote skewing by acting in different tissues. Conversely, *smax1* and *smxl2* could not suppress the *max2* waving phenotype individually but only in combination (Figure 5F, G), indicating that SMAX1 and SMXL2 act redundantly to promote waving. The role of SMAX1 and SMXL2 in the promotion of skewing and waving is consistent with the aforementioned role of KAI2 in inhibiting these processes. Intriguingly, we observed with plants grown in Munich [M] that *smxl678* was also able to suppress the *max2-1* skewing and waving phenotypes (Supplementary Figure 6A, B), consistent with the results of Swarbreck et al (2019). However, this was not the case in [L], where root skewing was often increased in *smxl678* relative to wild-type, and in which there was an additive increase in skewing in *smxl678 max2-1* (Supplementary Figure 6C). Notably, the effect of *kai2, smax1* and *smxl2* on skewing was consistent across all laboratories. It, therefore, seems that the KL repressors SMAX1 and SMXL2 promote root skewing constantly, while SMXL678 modulate root skewing in different directions probably dependent on other physiological parameters, which may be influenced by growth conditions. Thus, in the context of regulating root growth direction SMXL678 are likely not a direct target of KAI2.

**Figure 6.**
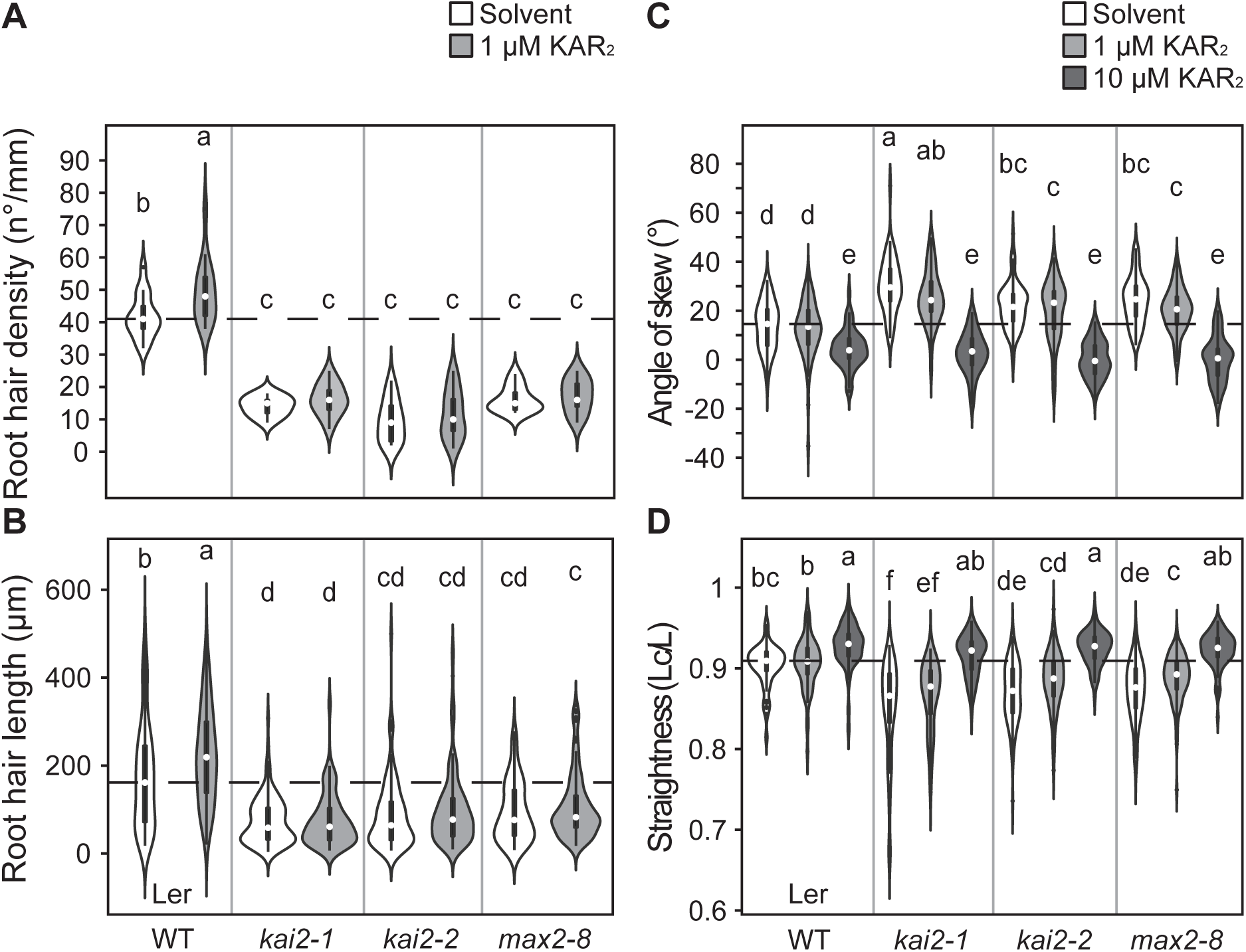
Effect of karrikin treatment on root growth and root hair development. **(A)** Root hair density, **(B)** root hair length, **(C)** root skewing and **(D)** root straightness of the indicated genotypes, treated with solvent (70% Methanol), 1 µM or 10 µM KAR_2_. The outline of the violin plot represents the probability of the kernel density. Black boxes represent interquartile ranges (IQR), with the white dot representing the median; whiskers extend to the highest and lowest data point but no more than ±1.5 times the IQR from the box; outliers are plotted individually. Different letters indicate different statistical groups (ANOVA, posthoc Tukey, **(A)** F_7,96_ = 60.79; p≤ 0.001, **(B)** F_7,975_ = 45.39; p≤ 0.001, **(C)** F_11,924_ = 90.19; p≤0.001, **(D)** F_11,924_ = 43.2; p≤0.001).

We also quantified the effect of SMXL mutations on root diameter. Interestingly, *smax1-2* alone could suppress the smaller root diameter of *max2-1* (Supplementary Figure 6D). In contrast to the skewing phenotype, the root diameter phenotype was not suppressed by *smxl2-1* alone or by *smxl678* in [M] (Supplementary Figure 6E, F). Thus, root diameter can be genetically separated from skewing, demonstrating that the two phenotypes are likely not causally related.

### Karrikin treatment enhances root hair development through KAI2

Having established unequivocal new roles of the karrikin receptor KAI2 in regulating root hair development, root growth direction and straightness, we focused on these phenotypes for further characterization and assessed if they can be influenced by exogenous addition of karrikin. We chose the Ler background because the Ler wild-type displays intrinsically stronger skewing than Col-0 (Figure 3, Supplementary Figure 3). Treatment with 1 µM KAR_2_ increased RHD and RHL relative to control treatments in a KAI2 and MAX2-dependent manner (Figure 6A, B), corroborating the role of KL-signalling in promoting root hair development.

Surprisingly, skewing and root straightness were not significantly influenced by 1µM KAR_2_., but 10μM KAR_2_ decreased root skewing and increased root straightness (Figure 6C, D). However, this high KAR_2_ concentration did affect skewing and straightness in all genotypes, including *kai2* and *max2*, suggesting non-specific side-effects of 10 µM KAR_2_ treatment (Figure 6C, D) (Swarbreck et al., 2019). Together this suggests that regulation of skewing and straightness in the wild-type is either saturated with endogenous KL and therefore cannot be significantly influenced by exogenous KAR_2_ or that the exogenously applied ligand is not reaching the cells, in which KAI2 needs to be triggered to suppress skewing.

### *rac*-GR24 enhances root hair development through both D14 and KAI2

Exogenous application of the strigolactone analogue *rac*-GR24 was previously shown to promote root hair elongation (Kapulnik et al., 2011b; Kapulnik et al., 2011a). We re-examined the effect of *rac*-GR24 on root hair development to understand whether it acts through perception by KAI2 or D14 (Scaffidi et al., 2014). Similar to KAR_2_, we found that *rac*-GR24 treatment increased both RHD and RHL (Figure 7A, B). This effect was dependent on *MAX2* as previously reported (Kapulnik et al., 2011b; Kapulnik et al., 2011a) but was independent of *KAI2*, suggesting that *rac*-GR24 might promote RHD and RHL through activation of the strigolactone receptor D14. We assessed this in detail and quantified RHD and RHL after treatment with the purified stereoisomers GR24^5DS^ (+GR24) and GR24^*ent*-5DS^ (-GR24), which are thought to activate D14 and KAI2, respectively (Scaffidi et al., 2014). We observed that both +GR24 and –GR24 promote RDH and RHL in the wild-type – although +GR24 is more active than –GR24, and both stereoisomers contained in *rac*-G24 appear to act additively (Supplementary Figure 7). Intriguingly, *d14* and *kai2* mutants respond in an unexpected manner to the stereoisomers: For *d14*, only -GR24 promotes RHD (as expected), but both -GR24 and +GR24 promote RHL to a similar degree (Supplementary Figure 7). Furthermore, *kai2-2* responded to both stereoisomers with both increased RHD and RHL, although +GR24 had a significantly stronger effect than -GR24 (Supplementary Figure 7). None of the stereoisomers promotes RHD and RHL in the *d14-1 kai2-2* double and *max2-1* mutants (Supplementary Figure 7). The first major implication of these results is that D14 can act to promote root hair development, when stimulated with ligand, even if that is not a standard endogenous function of D14 (Figure 1C). These results reconcile previous work that suggested that SL molecules can promote root hair development (Kapulnik et al., 2011b; Kapulnik et al., 2011a), with our results presented above that show SL signalling is dispensable for the regulation of root hair development. This is very similar to the hypocotyl, where D14-mediated SL perception can regulate hypocotyl elongation, but it not actually required to do so (Waters et al., 2012a; Scaffidi et al., 2013). The second major implication is that contrary to previously published data(Scaffidi et al., 2014) D14 perceives -GR24 ligands when KAI2 is absent, and KAI2 can perceive +GR24 ligands when D14 is absent, at least in the context of root hair elongation.

**Figure 7.**
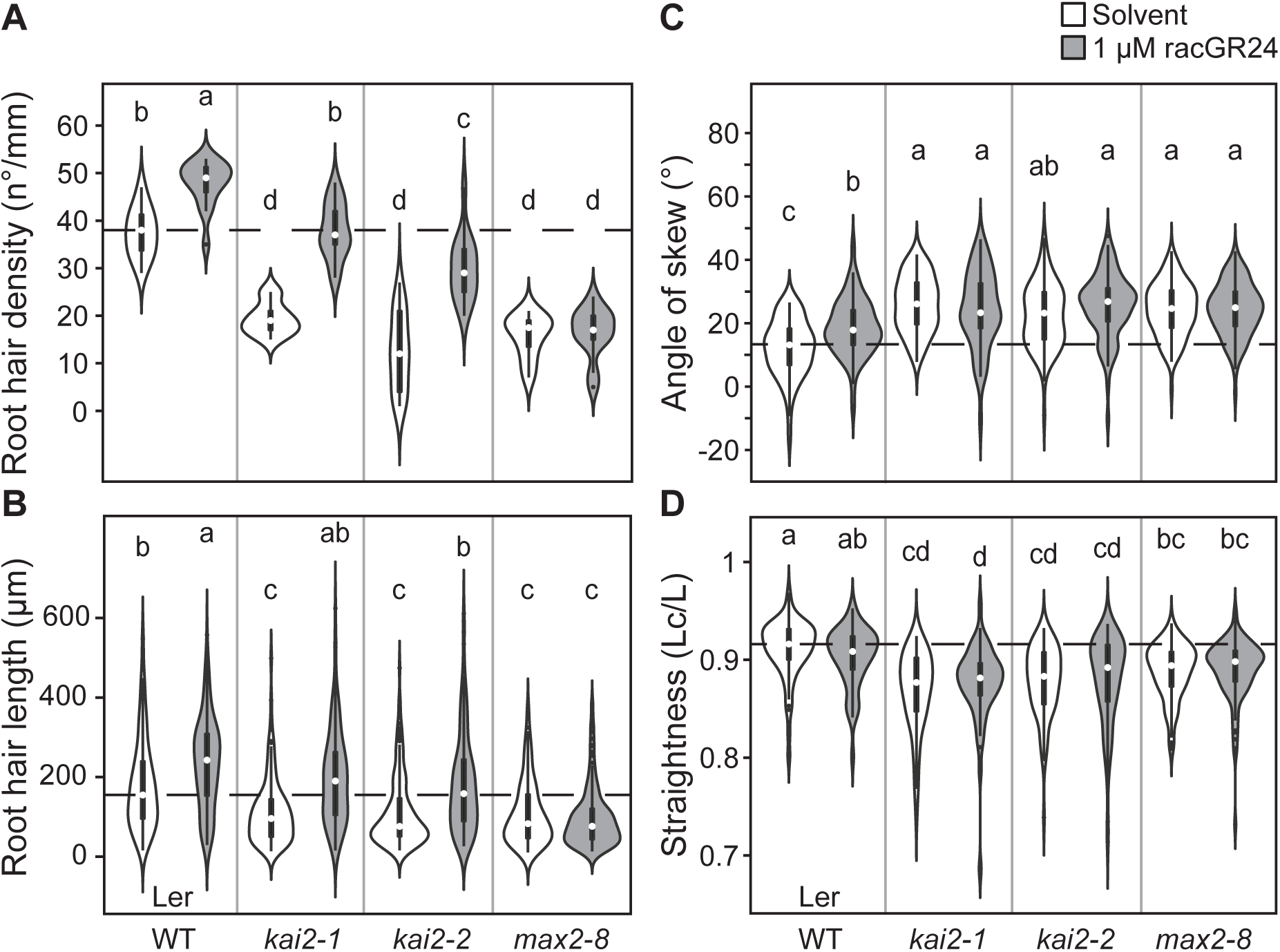
Effect of *rac*-GR24 on root growth and root hair development. **(A)** Root hair density, **(B)** root hair length, **(C)** root skewing and **(D)** root straightness of the indicated genotypes, treated with solvent (acetone) or 1 µM µM *rac*-GR24. The outline of the violin plot represents the probability of the kernel density. Black boxes represent interquartile ranges (IQR), with the white dot representing the median; whiskers extend to the highest and lowest data point but no more than ±1.5 times the IQR from the box; outliers are plotted individually. Different letters indicate different statistical groups (ANOVA, posthoc Tukey, **(A)** F_7,96_ = 60.79; p≤ 0.001, **(B)** F_7,1241_ = 39.81; p≤ 0.001, **(C)** F_7,624_ = 15.63; p≤0.001, **(D)** F_7,624_ = 10.73; p≤0.001).

We also examined the effect of *rac*-GR24 treatment on skewing and root straightness. Root straightness is not affected by *rac*-GR24 in any of the genotypes. Surprisingly, 1 µM *rac*-GR24 increases skewing in the wild-type to the level of *kai2* and *max2* mutants, while similar to KAR_2_ treatment none of the mutants respond to *rac*-GR24 (Figure 7C). This counter-intuitive response to *rac*-GR24 treatment may result from stimulation of D14 by the GR24 ligand to interact with MAX2, thereby competing with KAI2 for MAX2 and causing a *kai2*-like phenotype.

### KAI2 promotes root hair development by promoting auxin signalling

We decided to focus on root hair development as a model to understand the effects of KAI2 signalling, as this constitutes the most robust and easily observable phenotype. An extensive body of work shows that both root hair density and root hair elongation are regulated by auxin (Lee and Cho, 2008), so we postulated that KAI2 might regulate root hair development by modulating auxin signalling.

To examine this, we quantified the transcript accumulation of genes involved in root hair development, and auxin signalling, biosynthesis and transport in the *kai2-2* and *max2-8* mutants (Ler background) in comparison to the wild type. We chose an early time-point at 5 days post germination, when root hairs are already developing but lateral roots have not yet emerged, to avoid confounding effects caused by differences in lateral root numbers among the genotypes. We examined the expression of genes encoding the auxin biosynthesis and catabolism enzymes TAA1, YUC3, YUC6, DAO1, DAO2, auxin transporters AUX1, PIN2, PIN3, PIN7 (Cho et al., 2007; Lee and Cho, 2008; Ganguly et al., 2010), the auxin receptor TIR1 (Dharmasiri et al., 2005; Ganguly et al., 2010), and the auxin response factors ARF5, 7, 8, 19 (Mangano et al., 2017; Bhosale et al., 2018). Transcripts of all auxin biosynthesis, catabolism and transporter genes, *TIR1, ARF5* and *ARF8* accumulated to similar levels in all genotypes (Figure 8A, Supplementary Figure 8A). *ARF7* and *19* transcripts were significantly reduced in *kai2-2* and *max2-8*, similar to the well-established KAI2-response gene *DLK2* (Figure 8A, Supplementary Figure 8A). We also examined transcript levels of the bHLH transcription factor genes *RSL2, RLS4* and the expansin *EXP7*, which are all involved in root hair elongation and are likely induced downstream of the ARFs; as well as the phosphatidylinositol transfer protein-encoding gene *COW1* involved in root hair initiation (Cho and Cosgrove, 2002; Böhme et al., 2004; Yi et al., 2010; Pires et al., 2013). The expression of all four genes was significantly reduced in the KL perception mutants (Figure 8A). These results are therefore consistent with *kai2* root hair phenotypes arising through reduced auxin-mediated transcription in trichoblasts.

**Figure 8.**
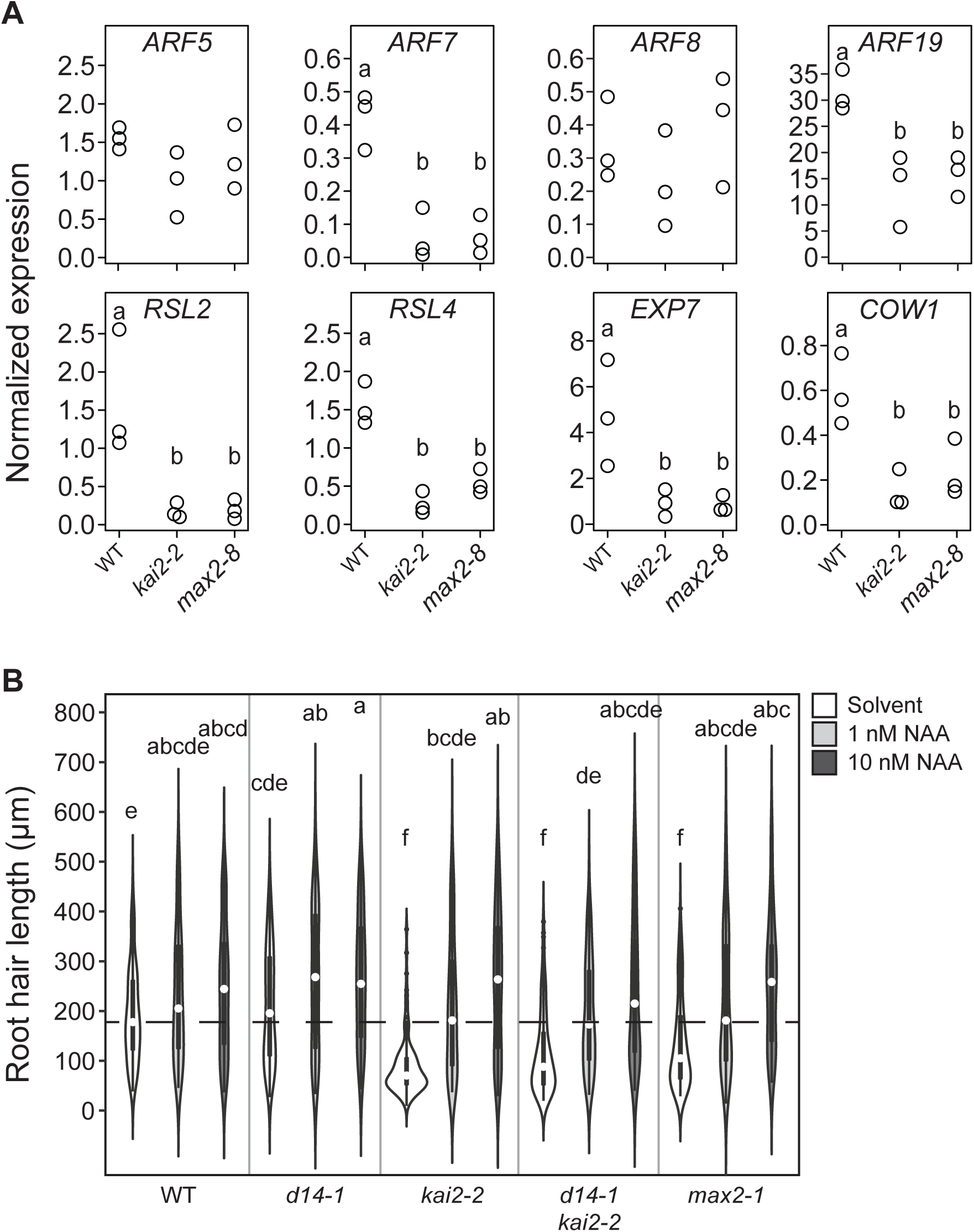
KL perception mutants alter auxin-inducible root hair transcriptional networks. **(A)** Transcript accumulation of *ARF5, ARF7, ARF8, ARF19, RSL2, RSL4, EXP7* and *COW1* in roots of the indicated genotypes. Expression levels of 3 biological replicates are normalized against those of *EF1α*. **(B)** Root hair length of the indicated genotypes, treated with solvent (96% Ethanol), 1 nM NAA or 10 nM NAA. The outline of the violin plot represents the probability of the kernel density. Black boxes represent interquartile ranges (IQR), with the white dot representing the median; whiskers extend to the highest and lowest data point but no more than ±1.5 times the IQR from the box; outliers are plotted individually. Different letters indicate different statistical groups (ANOVA, posthoc Tukey, **(A)** F_2,6_ = 0.251; p≤0.05 (*ARF5*), F_2,6_ = 31.69; p≤0.001 (*ARF7*), F_2,6_ = 0.03074; p≤0.05 (*ARF8*), F_2,6_ = 10.99; p≤0.001 (*ARF19*), F_2,6_ = 8.827; p≤0.001 (*RSL2*), F_2,6_ = 32.31; p≤0.001 (*RSL4*), F_2,6_ = 7.641; p≤0.05 (*EXP7*), F_2,6_ = 10.04; p≤0.05 (*COW1*), **(B)** F_7,1812_ = 25.83; p≤0.001).

To investigate whether *kai2* and *max2* roots are impaired in auxin signalling, we quantified the effect of mild (1 and 10nM) treatment with the auxin analogue NAA on various developmental parameters. Treatment with 1nM and 10nM NAA increases RHD of *kai2, d14 kai2* and *max2* mutants in a dose-dependent manner, although not sufficiently to completely restore RHD to the level of WT (Supplementary Figure 8B). However, only 1nM NAA is sufficient to completely restore RHL of KL perception mutants (Figure 8B), suggesting that *kai2* and *max2* are not impaired in auxin signalling *per se* but may have reduced auxin levels in certain cells types, in which auxin is required for root hair development. Consistent with this, both mutants show increased LRD in response to NAA treatment in a similar manner as the wild-type, indicating that they are indeed normally responsive to auxin (Supplementary Figure 8C).

### KAI2 alters longitudinal auxin distribution within the root system

Our data show that auxin signalling is reduced in *kai2* root hairs, but that *kai2* has no deficiency in responding to exogenous auxin. This suggests that *kai2* mutants either have reduced auxin levels, or altered auxin distribution within the root system. To distinguish between these possibilities, we determined the endogenous levels of indole-acetic acid (IAA) in the roots of *kai2-2* relative to WT and *d14-1, smax1 smxl2* and *smxl6,7,8*. Contrary to our expectations, we found that *kai2-2*, along with *d14-1*, had increased IAA levels relative to wild type, while IAA levels in *smax1 smxl2* and *smxl678* were similar to the wild type (Figure 9A). These elevated auxin levels are consistent with the increased LRD observed in both KL and SL perception mutants, but inconsistent with the root hair phenotypes observed in KL perception mutants, and their (partial) rescue by auxin treatment. Furthermore, given the similar auxin levels in *d14-1* and *kai2-2*, altered auxin levels in the whole root system are clearly not alone sufficient to explain the distinct pattern of root development in *kai2*. Given the increased LRD in the shootward part of *kai2* roots, and the reduced RHD/RHL at the root tip, we hypothesized that *kai2* may suffer from an altered longitudinal distribution of auxin within the root system, with a particular reduction in the root tip.

**Figure 9.**
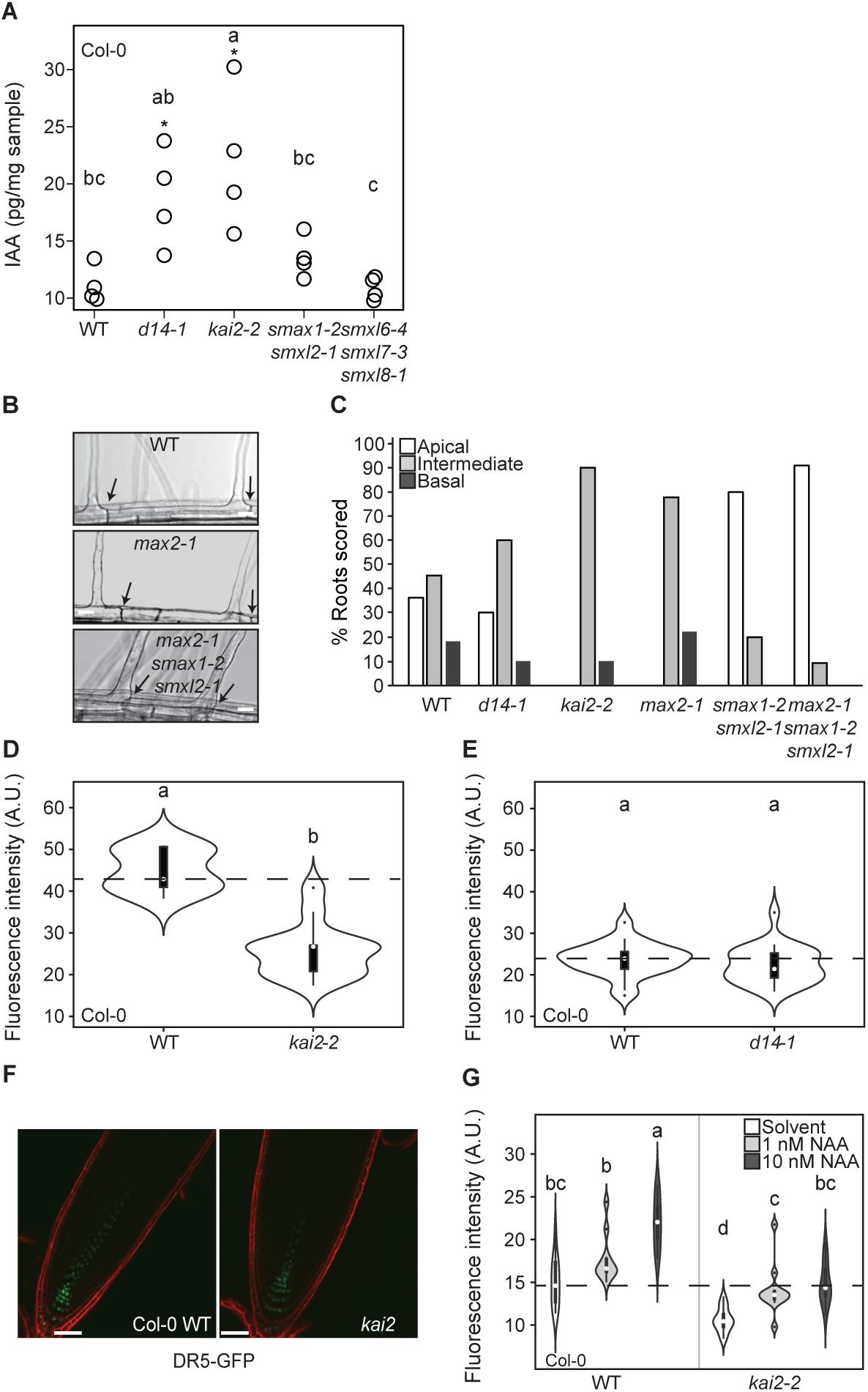
KL signalling alters longitudinal auxin distribution within the root system. **(A)** Measurement of free IAA (pg/mg sample) in 6dpg Arabidopsis roots of Col-0, *d14-1, kai2-* 2, *smax1-2 smxl2-1* and *smxl6-4 smxl7-3 smxl8-1*. **(B)** Images of representative trichoblasts showing sites of root hair emergence. Arrows indicate the most apical end of the cell. **(C)** Frequency distribution of different root hair positions observed in the indicated genotypes. **(D, E)** Fluorescence intensity (arbitrary units, A.U.) of *DR5v2:GFP* in the meristem zone of the indicated genotypes. **(F)** Confocal images of representative root tips of Col-0 wild type and *kai2-2* expressing the auxin reporter *DR5v2:GFP*, using the same microscope settings. **(G)** Fluorescence intensity (A.U.) of *DR5v2:GFP* in the meristem zone of Col-0 and *kai2-2* treated with solvent (96% Ethanol), 1 nM NAA or 10 nM NAA. The outline of the violin plot represents the probability of the kernel density. Black boxes represent interquartile ranges (IQR), with the white dot representing the median; whiskers extend to the highest and lowest data point but no more than ±1.5 times the IQR from the box; outliers are plotted individually. Different letters indicate different statistical groups (ANOVA, posthoc Tukey, **(A)** F_4,15_ = 7.544; p≤ 0.01, **(D)** F_1,21_ = 60.06; p≤0.001, **(E)** F_1,45_ = 1.239; p≤0.05, **(G)** F_5,67_ = 24.69; p≤0.001). Asterisks indicate a significant difference from wild type (Student’s t-test, p≤0.05).

The positioning of tip growth along the longitudinal axis of root hair cells has been shown to be sensitive to an auxin gradient generated in the root tip (Fischer et al., 2006). We reasoned that if *kai2* mutants indeed have reduced auxin levels in the root tip, we would see altered root hair positioning in addition to reduced density and elongation. In wild-type and *d14* mutants, root hairs are approximately equally distributed between basal (rootward) and intermediate positions, but in contrast, most root hairs of *kai2* and *max2* are shifted towards an intermediate position (Figure 9B, C). Conversely, for *smax1 smxl2* and *max2 smax1 smxl2* mutants, a much higher proportion of root hairs than in wild type were positioned basally (Figure 9B, C). These data are thus consistent with the root hair phenotypes in KL signalling mutants arising from reduced auxin levels in the root tip.

To confirm this idea, we examined the expression of the highly auxin-sensitive *DR5v2:GFP* reporter (Liao et al., 2015) in the root meristem of *kai2-2* mutants, and observed a marked reduction in *DR5v2:GFP* expression as compared to wild type (Figure 9D, F). This difference in *DR5v2:GFP* expression was not observed in the *d14-1* mutant background (Figure 9E). Consistent with previous results, we found that treatment with 1nM NAA is sufficient to restore *DR5v2:GFP* expression to wild type levels in *kai2-2*. The dose-response pattern of *kai2-2* to NAA is comparable to wild-type in this system, again confirming there is no lack of auxin sensitivity in *kai2-2* (Figure 9G). In summary, our data indicate that *kai2-2* has globally increased auxin levels within the root system, but that this auxin is incorrectly distributed along the longitudinal axis of the root, resulting in a reduction in auxin levels in the root meristem.

### KAI2 regulates PIN7 abundance in the differentiation zone

In the shoot system, strigolactones regulate the abundance of PIN1 protein and thereby control auxin transport along shoot axes (Shinohara et al., 2013). To explain the altered longitudinal distribution of auxin within the root system, we hypothesized that KAI2 might also regulate PIN protein abundance in the root and examined GFP-fusion of PIN1, 2, 3, 4, and 7 in *kai2* roots. We could not detect significant *PIN1:PIN1-GFP, PIN2:PIN2-GFP* or *PIN4:PIN4:GFP* expression outside the meristem zone in either *kai2* or wild type, but we observed strong expression of *PIN3:PIN3-GFP* and *PIN7:PIN7-GFP* along the whole root axis in both wild type and *kai2-2* (Figure 10A). We did not observe changes in PIN3-GFP abundance in *kai2-2* (Figure 10E, F), but intriguingly, PIN7-GFP abundance in the stele is increased in *kai2-2* relative to wild type in the younger parts of the differentiation zone (Figure 10I, J). However, in the older parts of the differentiation zone and in the meristem zone, there was only a minor difference in PIN7-GFP abundance in the stele, which was only statistically significant when a T-test instead of ANOVA (comparing data for all zones) was used (Figure 10I). Publicly available gene expression data show that KAI2 is strongly expressed in the stele in the elongation and differentiation zones of the root, but not in the meristem zone (Brady et al, 2007), which is consistent with the observed location of changes in PIN7 abundance (Supplementary Figure 8A). We observed that *PIN7* mRNA accumulation is not increased in *kai2-2* roots (Supplementary Figure 8A). Thus, we propose that, analogously to the effect of strigolactones on PIN1, KAI2 signalling negatively regulates PIN7 at the post-transcriptional level. Alternatively, an increase in *PIN7* transcript accumulation in a small root zone is not visible in cDNA from entire roots.

**Figure 10.**
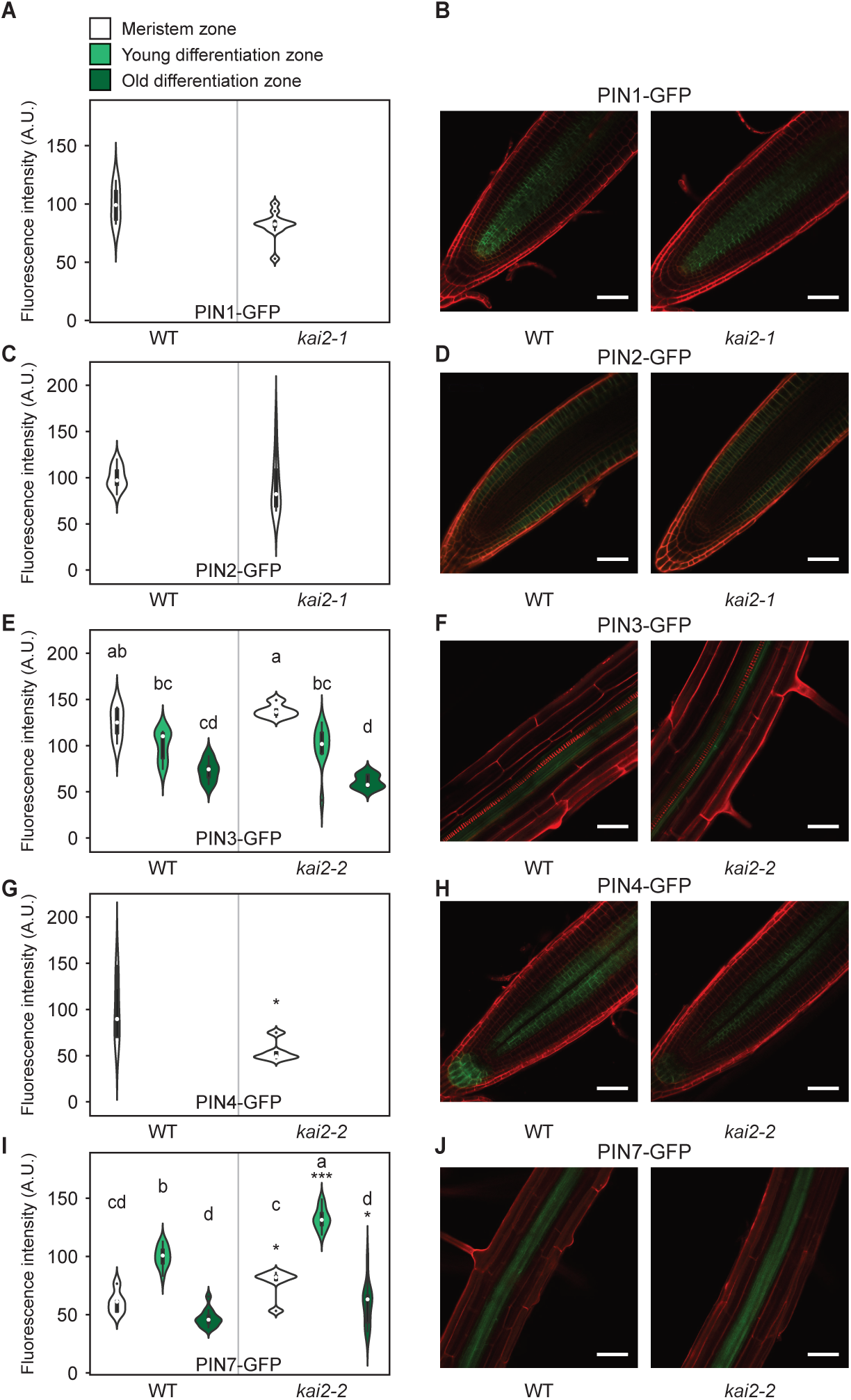
Accumulation of PIN4- and PIN7-GFP is altered in KAR perception mutants. Fluorescence intensity (arbitrary units, A.U.) of PIN1-GFP **(A)**, PIN2-GFP **(C)**, PIN3-GFP **(E)**, PIN4-GFP **(G)**, PIN7-GFP **(I)**, in Col-0 wild type and *kai2-2* in the meristem zone, young differentiation zone and old differentiation zone (see materials and methods for a description of the zones). **(D)** Confocal images of representative roots of Col-0 wild type and *kai2-1* or *kai2-2* expressing PIN1-GFP in the meristem zone **(B)**, PIN2-GFP in the meristem zone **(D)**, PIN3-GFP in the young differentiation zone **(F)**, PIN4-GFP in the meristem zone **(H)** and PIN7-GFP in the young differentiation zone **(J)** shown as green fluorescence. Roots were counterstained with propidium iodide (red fluorescence). Scale bars, 50 µm. The outline of the violin plot in **(A)**, **(C)**, **(E)**, **(G)**, **(I)** represents the probability of the kernel density. Black boxes represent interquartile ranges (IQR), with the white dot representing the median; whiskers extend to the highest and lowest data point but no more than ±1.5 times the IQR from the box; outliers are plotted individually. Different letters indicate different statistical groups (ANOVA, posthoc Tukey, **(E)** F_5,36_ = 14.97; p<0.001, **(I)** F_5,53_ = 88.22 p≤0.001,). Asterisks indicate a significant difference from wild type (Student’s t-test, ***p≤0.001, *p≤0.05).

## DISCUSSION

Root systems flexibly adapt their architecture and morphology to hetereogenous soil environments and to the physiological needs of the plant. A network of plant hormone signalling pathways is essential for translating environmental signals and physiological states into developmental outputs (Osmont et al., 2007). Strigolactones (SLs) have been assumed to play an important role in modulating root development (Kapulnik et al., 2011b; Ruyter-Spira et al., 2011; Jiang et al., 2015). However, the frequent use of *max2* mutants and the synthetic ‘SL analogue’ *rac*-GR24 have led to ambiguous assignment of roles of SLs to lateral root and root hair development, because MAX2 and *rac*-GR24 also participate in karrikin-like (KL) signalling (discussed in Waters et al. 2017). Using a collection of mutants specifically defective in either the SL or KL pathway, we have carefully dissected the role of the two signalling modules in root and root hair development, and overall our data indicate that in Arabidopsis seedlings KL signalling plays a much larger role in root development than SL signalling.

### KL signalling regulates lateral root density together with SL signalling

We observed that LRD is consistently higher in *kai2* mutants than wild type, indicating that KL signalling plays a role in modulating lateral root numbers. This is consistent with previous reports showing increased LRD in *max2* and suppression of lateral root emergence by *rac*-GR24 (Jiang et al., 2015). In our study, SL biosynthesis and perception mutants also display subtle changes in root architectural parameters, such as primary root length (PRL) and LRD. However, root architecture phenotypes of *d14* and SL biosynthesis mutants were much less reproducible than the *kai2* phenotypes and varied among experiments and locations (Leeds [L] vs Cambridge [C]), indicating that the role of SL signalling in the regulation of PRL and LRD strongly depends on environmental conditions. In [L], the PRL phenotype previously described by Ruyter-Spira et al. (2011), was observed, but not the LRD phenotype (Jiang et al., 2015) and in [C] the opposite was true. This suggests that these phenotypes are to some extent mutually exclusive, and that expression of one phenotype reduces expression of the other, which may explain some of the contradictory reports regarding effects of SLs on root development (Ruyter-Spira et al., 2011; Jiang et al., 2015). Nevertheless, the double *d14 kai2* mutant showed the largest increase in LRD indicating that both signalling pathways contribute to modulating LRD. This is confirmed by suppression of the *max2* LRD phenotype by mutants in the targets of both KL (SMAX1/SMXL2) and SL (SMXL678) signalling (see below).

### KL signalling suppresses skewing and waving

No single signalling pathway for control of root skewing and straightness has been identified, but several studies expose different pathways impinging on these root behaviours (reviewed in Roy and Bassham. 2014). The activities of multiple hormones, such as auxin and ethylene, are among the candidates (Buer et al., 2003; Qi and Zheng, 2013). Here we demonstrate that KL signalling is a novel regulator of root skewing and root straightness. The increased skewing and waving phenotype of KL perception mutants were found in both the Col-0 and Ler background although Ler shows an intrinsically higher tendency to skew than the wild type. Our results are broadly consistent with the recent report of Swarbreck et al., (2019), but our interpretation of the phenotypes differs. Swarbreck et al., (2019), speculate that skewing may be caused by increased epidermal cell file rotation and smaller root diameter of *kai2* mutants. We did not observe epidermal cell file rotation but rather shorter epidermal cells. Since not only *kai2* but also *d14* have a reduced epidermal cell length we conclude that this is not related to skewing. In addition, we propose that the reduced root diameter of KL perception mutants is not causative for skewing because *sxml678* could suppress skewing of *max2* in [M] but did not concomitantly suppress the decreased root diameter.

Interestingly, although exogenous treatment with KAR_2_ led to an increase in RHD and RHL we were unable to reduce skewing of Ler wild type with karrikin treatment in a specific manner, and this was also reported by Swarbreck et al., (2019). It is possible that for skewing, the system is already saturated with endogenous KL, at least in the applied growth conditions. Alternatively, exogenously applied karrikin or a possible breakdown product (Waters et al., 2015), may not reach the cells in which in it needed to affect skewing.

### KL signalling is a key regulator of root hair development

One of our major findings is the strong impact of KL signalling on root hair development. *kai2* and *max2* mutants in both Col-0 and Ler backgrounds show clear defects in RHD, RHL and root hair positioning along the longitudinal axis of trichoblasts. These root hair phenotypes also consistently show the opposite phenotype in *smax1 smxl2* mutants, and in addition, karrikin treatment of wild type increases RHD and RHL. Our results thus present compelling evidence that KL signalling is a key regulator of root hair development.

The two stereoisomers of *rac*-GR24, *5DS*-GR24 (+GR24) and *ent5DS*-GR24 (-GR24) have been suggested to specifically activate D14 and KAI2, respectively (Scaffidi et al., 2014). Our work suggests that the perception and response of these GR24 stereoisomers may not be completely specific, at least in the case of root hair development. The *kai2* mutant responds with increased RHD to both +GR24 and -GR24, while surprisingly, for RHL both the *d14* and the *kai2* mutant respond to both +GR24 and -GR24. Although it has been shown by differential scanning fluorimetry (DSF) *in vitro* that KAI2 binds only -GR24, while D14 showed binding of both +GR24 and -GR24 (Waters et al., 2015) the situation *in vivo* may be different and binding of both ligands to both α/β hydrolase receptors D14 and KAI2 may be stabilized through receptor protein complexes. Although binding of the ‘wrong’ stereoisomer to the α/β hydrolase receptor may be less efficient than binding of the ‘correct’ one it may suffice to trigger developmental responses which are very sensitive to removal of SMXL proteins – which may be the case for RHL - or which may require other additional interaction partners in the receptor complex than others.

Despite the possibility to trigger RHD and RHL with additional ligand in the *kai2* mutant background, mutations of *D14* or SL biosynthesis genes did not have an impact on root hair development in our study. (Mayzlish-Gati et al., 2012) found a small decrease in RHD of the SL biosynthesis mutant *max4-1*, which is inconsistent with our data. It is possible that RHD can be regulated via D14 under certain conditions, when SL levels are very high, for example under phosphate starvation (Ito et al., 2015). This assumption is supported by our results showing that *rac*-GR24 and both GR24 stereoisomers can trigger an increase in RHD and RHL in *kai2* but not in *max2* mutants, demonstrating that D14 participates in the promotion of RHD when additional ligand is supplied. Nevertheless, KAI2 instead of D14 being the major regulator of root hair development seems to make sense also from an evolutionary point of view. Root hair development and tip growth in Arabidopsis rely on conserved functions and genes, which also operate in the development of rhizoids of *Marchantia polymorpha* gametophytes, which appear to be homologous to root hairs (Tam et al., 2015; Honkanen and Dolan, 2016; Honkanen et al., 2016). D14 occurs only in the seed plants and while KAI2 is already present in algae (Delaux et al., 2012; Waters et al., 2012b; Bythell-Douglas et al., 2017). It is thus conceivable that KAI2-mediated signalling may be part of an ancient and conserved pathway regulating tip growth of epidermal cells.

### SL and KL signalling in the root are canonical in nature

We have previously highlighted some phenotypic characteristics suggesting that SL and KL signalling in the root might not act through the canonical D14-SMXL678 and KAI2-SMAX1 receptor-repressor pairs (Waters et al., 2017). Reasons for this suggestion were a) that *max2* mutants had stronger LRD phenotypes than SL biosynthesis mutants (Kapulnik et al., 2011b; Ruyter-Spira et al., 2011; Jiang et al., 2015), which increased the likelihood that KAI2 regulates lateral root emergence rather than or in addition to D14, and b) that mutations of the canonical SL signalling repressor genes *SMXL678* was able to completely suppress the *max2* LRD phenotype, while *smax1* was unable to do so (Soundappan et al., 2015; Waters et al., 2017). We now show that *smax1* in combination with a mutation in the partially redundant *SMXL2* (Stanga et al., 2016) can suppress the LRD phenotype of *max2*, demonstrating that the canonical KL signalling repressors are involved in regulating lateral root formation and that *SMXL2* compensates for the absence of functional *SMAX1* in lateral root development (Soundappan et al., 2015). The suppression of the *max2* LRD phenotype by *smxl678* as well as *smax1 smlx2* is consistent with our observation that both D14 and KAI2 regulate LRD. Although we confirm that *smxl678* completely suppresses *max2* in terms of absolute phenotype, it is not completely epistatic to *max2*, and the KL-related part of the *max2* LRD phenotype remains unsuppressed in *smxl678 max2*. The same is true for *smax1 smxl2*, which completely suppress *max2* LRD phenotypes in absolute terms, but are not completely epistatic to *max2*. Thus, the accumulation of both SMAX1/SMXL2 and SMXL678 contributes to *max2* LRD phenotypes and there is no evidence to suggest these responses are mediated by non-canonical receptor-repressor pairs.

The case is even more clear-cut for RHL, RHD, root straightness and root diameter, for which only KL perception mutants show a phenotype and this is suppressed by mutating *SMAX1* and *SMXL2*. In the case of root diameter, mutation of *SMAX1* is sufficient to suppress the *max2* phenotypes (Supplemental Table 1). This partial redundancy of SMAX1 and SMXL2 is also seen in seed germination, hypocotyl growth and leaf shape (Soundappan et al., 2015; Stanga et al., 2016). This likely arises from different expression patterns of the two genes: in tissues where only one of the two proteins is expressed, removing this one is sufficient to suppress the phenotype. Conversely, in the case of skewing, removing either SMAX1 or SMXL2 is sufficient to suppress the *max2* phenotype (Table S1), suggesting that skewing is particularly sensitive to SMAX1/SMXL2 levels or stochiometry or that SMAX1 or SMXL2 regulate skewing in different tissues.

Swarbreck et al., (2019), suggested that KL signalling might act to degrade both SMAX1/SMXL2 and SMXL678 to regulate root skewing. Here we reject the idea that KL signalling regulates skewing through SMXL678 (Swarbreck et al., 2019). Although *smxl678* can suppress skewing of *max2* in some locations ([M]), in other locations ([L]) it increased skewing additively with *max2*. Thus, although SMXL678 can certainly regulate skewing, this appears to be independent of the defined effect of KL signalling. The location-dependent opposite skewing behaviour of *max2 smxl678* mutants suggests that the role of SMXL678 in skewing may strongly depend on environmental conditions, and it will be interesting to identify the mechanisms underlying this phenomenon in the future.

Ruyter-Spira et al., (2011), previously suggested that the impact of SLs on root development might be best understood as a reflection of their effect on the auxin landscape: assuming a bell-shaped auxin response, the same concentration of SL might lower the auxin levels from an optimal state to a sub-optimal level, or from a super-optimal level to an optimal level — and thus have opposite developmental effects depending on the background auxin concentration. Such as scenario might well underlie the strong sensitivity of SL root responses to growth conditions, since temperature and light both affect endogenous auxin levels (Gray et al., 1998; Ljung, 2013).

### KL signalling modulates auxin distribution in the root

We focused on root hair development to investigate downstream events of KL signalling in the root. In agreement with reduced RHD and RHL we hypothesized that auxin signalling may be affected. Indeed, transcript accumulation of the auxin response factor genes *ARF7* and *ARF19*, which have been implicated in controlling root hair elongation (Mangano et al., 2017; Bhosale et al., 2018), is reduced in *kai2* and *max2* mutants at 5dpg and before lateral roots have emerged. This is accompanied with a reduction in transcript accumulation of putative ARF7/ARF19 downstream target genes *RSL2, RSL4, EXP7* and *COW1* that are regulators and executors of root hair development (Böhme et al., 2004; Yi et al., 2010; Shibata et al., 2018). The reduced expression of *ARF7* and *19* in KL perception mutants coincides with a reduced planar polarity of root hairs and expression of the auxin reporter *DR5v2:GFP* in the primary root meristem of *kai2*. Together, this suggests that insufficient auxin in the *kai2* root meristem causes reduced expression of *ARF7* and *19* leading to defects in root hair development. Indeed, we can fully restore RHL and *DR5v2:GFP* expression in *kai2* with exogenous NAA treatment. RHD cannot be completely restored by exogenous NAA. This may indicate that, in the absence of functional KAI2, NAA – although it is sufficient to promote elongation of already initiated root hairs - is not correctly distributed within the root tissue and thus unable to induce specification of the same number of trichoblasts as in the wild type.

Since PIN proteins are important for auxin distribution in the Arabidopsis root meristem and differentiation zone (Blilou et al., 2005) and SL signalling has previously been shown to affect PIN1 abundance at plasma membranes in the shoot (Shinohara et al., 2013), we investigated whether KAI2 affects the expression or distribution of PIN1, 2, 3, 4, and 7 GFP-fusions expressed under the control of their endogenous promoters. While accumulation of most PIN proteins did not significantly change between wild type and *kai2*, PIN7-GFP accumulated to increased levels throughout the root, and especially in the young differentiation zone of the *kai2* mutant. This is consistent with the observation that *rac*-GR24 treatment, (which increases KAI2-mediated signalling) leads to reduced PIN7-GFP accumulation in the Arabidopsis root vasculature (Ruyter-Spira et al., 2011). PIN7 has been observed to localize to basal as well as lateral membranes in the stele (Blilou et al., 2005; Marhavý et al., 2013). We, therefore, speculate that increased PIN7 abundance in the stele leads to increased leakage of auxin into the surrounding tissue, thus reducing overall rootward auxin flux. This leakage into the outer tissues likely explains the increased lateral root emergence in *kai2* and *max2* mutants, which is an auxin-driven process (Péret et al., 2012; Vermeer et al., 2014). Thus, an aberrant longitudinal auxin distribution, which may at least partly depend on increased PIN7, may explain the *kai2* lateral root and root hair phenotypes, which are at first sight contradictory. In this context, it is interesting that endodermis-specific expression of *MAX2* driven by the *SCARECROW* promoter was found to be sufficient to rescue RHD and LRD in a *max2* mutant (Koren et al., 2013; Madmon et al., 2016), consistent with a non-cell autonomous signalling pathway. This, it is possible that the KAI2-MAX2 regulation of PIN7 in the endodermis is particularly important for controlling auxin distribution in the root.

Taking together our results, we propose the following model to explain the root and root hair developmental defects in *kai2* mutants (Figure 11). 1) PIN7 over-accumulates in the differentiation zone of *kai2* roots, leading to increased auxin leakage from the stele into the surrounding tissues. 2) This auxin promotes lateral root emergence. 3) Leakage of auxin reduces auxin delivery to the meristem zone. 4) Reduced auxin in the meristem zone causes shallower asymmetries in responses to gravity and touch stimuli, leading to altered growth vectors. 5) Reduced auxin distribution along the root epidermis reduces root hair planar polarity, density and elongation. Conversely, in *smax1 smxl2* mutants, there is likely decreased PIN7 abundance leading to reduced auxin leakage from the stele, reducing lateral root emergence, and increasing auxin delivery to the root meristem, thereby promoting straighter root growth, and increased polarization, density and elongation of root hairs (Figure 11).

**Figure 11.**
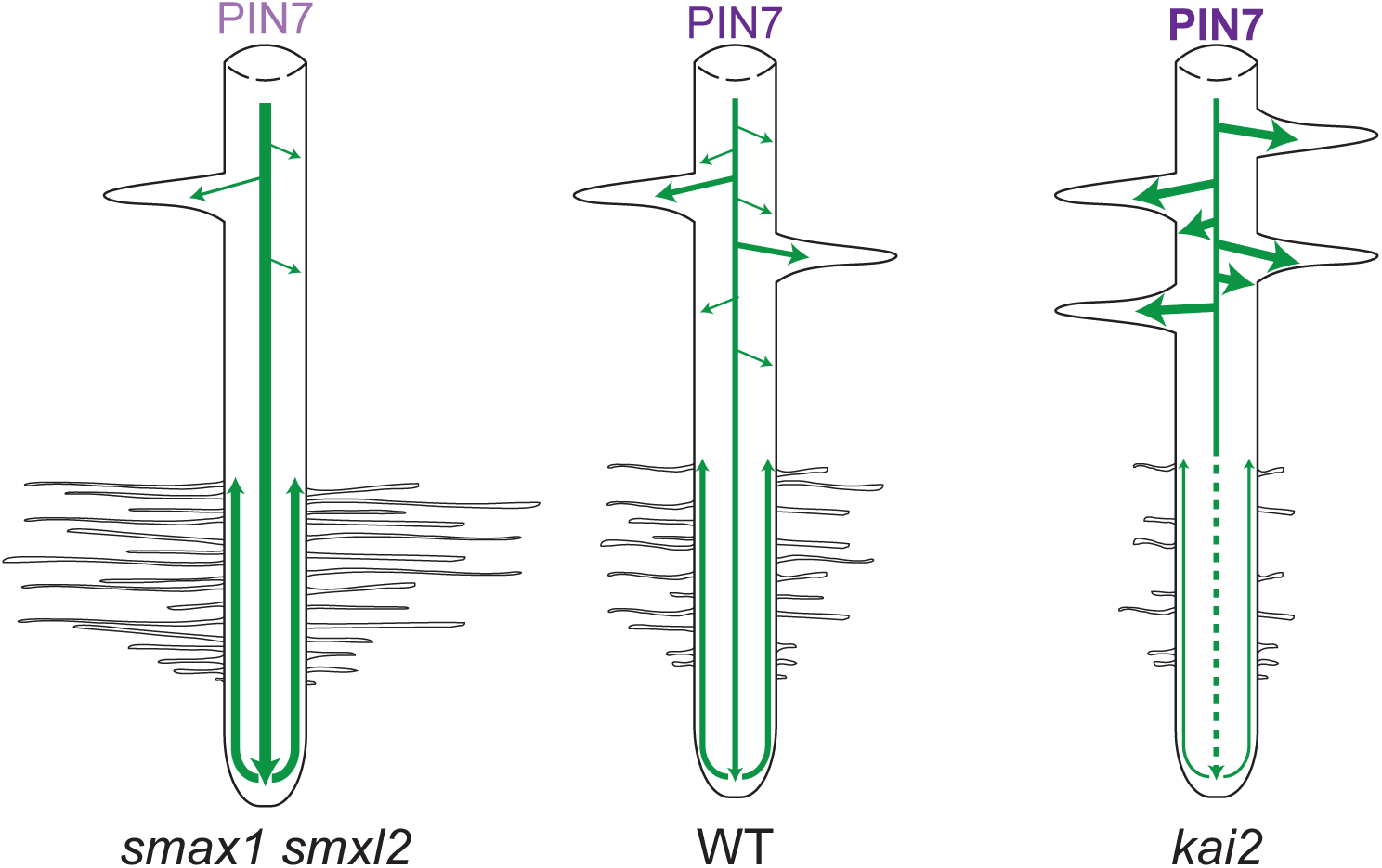
Model for auxin distribution in Arabidopsis roots. Hypothetical model showing auxin distribution (green) of *smax1 smxl2*, wild type and *kai2* roots. Root hair length and density is reduced in *kai2* but increased in *smax1smxl2* roots. Lateral root density is increased in *kai2* but reduced in *smax1 smxl2*. Bold purple text indicates over-accumulation of PIN7 in *kai2*, pink text indicates unknown accumulation of PIN7 in *smax1 smxl2*.

Overall our results strongly suggest that SMAX1 and SMXL2 play an important role in controlling root and root hair development by regulating the auxin landscape in the root. However, some traits such as LRD and epidermal cell length are regulated by both SMAX1/SMXL2 and SMXL678 and both *kai2* and *d14* mutants accumulate more auxin in their roots. Key challenges for future studies will be to understand how exactly SMXL proteins regulate PIN abundance and whether the pathways have additional downstream targets in the root, beyond the regulation of auxin distribution. We do not currently know enough about the upstream inputs into the KL signalling pathway to understand the aetiology of KAI2-induced root development, but undoubtedly the phenotypes described here will provide important clues and tools in this regard. SL production increases in several plant species upon phosphate starvation (López-Ráez et al., 2008; Yoneyama et al., 2013; Sun et al., 2014; Decker et al., 2017), but it is yet unknown whether KL signalling is also influenced by mineral nutrient levels. However, expression of KAI2 does respond to light conditions, and thus KL signalling could potentially integrate light cues into root development (Sun and Ni, 2011). Indeed, it is likely that both signalling pathways are influenced by multiple abiotic and perhaps biotic stimuli, and it will be exciting to learn how SL and KL signalling tune root development to environmental conditions.

## MATERIALS AND METHODS

### Plant material

*Arabidopsis thaliana* genotypes were in Columbia-0 (Col-0) or Landsberg *erecta* (Ler) parental backgrounds. The *max1-1, max2-1, max2-2* (Stirnberg et al., 2002), *max2-8* (Nelson et al., 2011), *max3-9* (Booker et al., 2005), *max4-5* (Bennett et al., 2016b), *d14-1* (Waters et al., 2012a), *kai2-1, kai2-2* (Nelson et al., 2011), *kai2-2* [Col-0], *d14-1 kai2-2* (Bennett et al., 2016a), *smax1-2, max2-1 smax1-2, smax1-3, max2-2 smax1-3* (Stanga et al., 2013), *smax1-2 smxl2-1, max2-1 smax1-2 smxl2-1* (Stanga et al., 2016), *smxl6-4 smxl7-3 smxl8-1, max2-1 smxl6-4 smxl7-3 smxl8-1* (Soundappan et al., 2015), *DR5v2:GFP* (Liao et al., 2015), *PIN3:PIN3-GFP* and *PIN7:PIN7-GFP* (Bennett et al., 2016b; Bennett et al., 2016a) lines have all been previously described.

### Plant growth conditions

For analysis of root growth, *Arabidopsis thaliana* seeds were grown in axenic conditions on 12×12cm square plates containing 60 ml agar-solidified medium. Seed were surface sterilized either by vapour sterilization, or by washing with 1 ml of 70% (v/v) ethanol and 0.05% (v/v) Triton X-100 with gentle mixing by inversion for 6 minutes at room temperature, followed by 1 wash with 96% ethanol and 5 washes with sterile distilled water. In Cambridge [C] (PRL, LRD) and Leeds [L] (measurements of PRL, LRD, DR5-GFP expression and PIN7-GFP expression), seedlings were grown on plates containing ATS medium (Wilson et al., 1990) supplemented with 1% sucrose (w/v) and solidified with 0.8% ATS. In Munich [M] (measurement of RHD, RHL, root hair position, skewing, waving, cell length and root diameter), seedlings were grown on plates containing 0.5X Murashige & Skoog medium, pH5.8 (½ MS) (Duchefa, Netherlands), supplemented with 1% sucrose and solidified with 1.5% agar. Plates were stratified at 4°C for 2-3 days in the dark, and then transferred to a growth cabinet under controlled conditions at 22 °C, 16-h/8-h light/dark cycle (intensity ∼120 µmol m^−2^ s^−1^). Unless otherwise indicated, plates were vertically-inclined

### Phytohormone treatments

NAA was purchased from Sigma-Aldrich (St. Louis, United States), *rac*-GR24 from Chiralix (Nijmegen, The Netherlands), *ent5DS*-GR24 (-GR24) and *5DS*-GR24 (+GR24) from Strigolab (Turin, Italy), and KAR_2_ from Olchemim (Olomouc, Czech Republic). NAA was dissolved in either 2% DMSO, 70% ethanol for a 1mM stock, or 100% ethanol for the preparation of 10 mM stock solution. For treatment with *rac*-GR24, *ent5DS*-GR24 or *5DS*-GR24, 1 mM stock solutions were prepared in 100% acetone. KAR was dissolved in 70% methanol for the preparation of 1 mM stock. The volume required to reach the final concentration of these different stock solutions was added to molten media prior to pouring Petri dishes. In each experiment, an equivalent volume of solvent was added to Petri dishes for untreated controls.

### Primary and lateral root quantification

For quantification of primary root length and lateral root number, seedlings were grown as described above in [C] and [L] for 10 days after germination (dag), except where stated. This allowed for emergence of lateral roots sufficient for quantification in wild-type seedlings. A dissecting microscope was used to count emerged lateral roots in each root system, and images of the plates were then taken using a flatbed scanner. Primary root length was quantified using Image J. Separate experiments were primarily used to assess root skewing (see below), but root skewing angles were also measured from these images generated in these experiments.

### Root skewing and straightness assay

The root slanting assay was modified from the method described by Rutherford and Masson (1996). Arabidopsis seedlings were grown in [M] under the conditions described above (except for Supplementary Figure 6C for which plants were grown in [L]). Images were taken 5 days after germination using an Epson Perfection V800 Pro Scanner. Images were analysed using the Simple Neurite Tracer plug-in of Fiji (https://imagej.net/Fiji/Downloads) to determine the following parameters as illustrated in Figure 3; root length (L), ratio of the straight line between the hypocotyl-root junction and the root tip (Lc), and vertical axis (Ly). These measurements were used to calculate the root skewing angle (α) and root straightness (Lc/L) as previously described (Grabov et al., 2005; Vaughn and Masson, 2011).

### Determination of root hair density, length and position

Root hair growth was examined in [M] on the same *Arabidopsis* roots, which were used for determining root skewing and straightness. Images were taken at 2 mm from the root tip of a minimum of 10 roots per genotype and treatment with a Leica DM6 B microscope equipped with a Leica DFC9000 GT camera. The number of root hairs was determined by counting the root hairs between 2 and 3 mm from the root tip on each root, and root hair length was measured for 10 −15 different root hairs per root using Fiji. The root hair position was determined following the method described by (Masucci and Schiefelbein, 1994) for 5-15 root hairs per root and a minimum of 10 roots per genotype.

### Root diameter and cell length analysis

Using the same images as for root hair quantification, we analyzed root diameter, root cell length and number of cells using Fiji. Root diameter was measured at 2.5 mm from the root tip. The number of cells was defined as the number of epidermal cells that crossed a 1-mm-long straight line drawn between 2 to 3 mm from the root tip. Root cell length was measured for at least 10 different epidermal cells per individual root, between 2 to 3 mm from the root tip.

### Confocal microscopy

Laser-scanning confocal microscopy was performed on either Zeiss LSM700 or LSM880 imaging system with a 20X lens. Roots were stained with propidium iodide (10ug/ml) and mounted on slides. GFP excitation was performed using a 488 nm laser, and fluorescence was detected between 488 and 555nm. Propidium iodide excitation was performed using a 561 nm laser, and fluorescence was detected between above 610nm. The same detection settings were used for all images captured in a single experiment. GFP quantification was performed on non-saturated images, using Zeiss ‘ZEN’ software. For *DR5v2:GFP* expression, this was done on a region of interest that included all columella and quiescent centre nuclei. For PIN7-GFP, fluorescence was quantified in a region of interest covering the stele of the root.

### Auxin quantification

Seedlings were grown for 6 days on ATS-agar medium with sucrose, then the roots were dissected from the seedlings in pools of ∼30 roots. From these samples (10-20 mg fresh weight) IAA was purified and analyzed by gas chromatography-tandem mass spectrometry (GC-MS/MS) as described (Andersen et al., 2008) with minor modifications. To each sample, 500 pg ^13^C_6_-IAA was added as an internal standard before extraction. Four replicates were analyzed for each genotype.

### RNA extraction and gene expression analysis

For analysis of transcript levels by qRT-PCR, a minimum of 100 roots per sample was rapidly shock frozen in liquid nitrogen. RNA was isolated using NucleoSpin RNA plant and fungi kit (Macherey-Nagel). The concentration and purity of RNA were evaluated with DS-11 FX+ spectrophotometer/fluorometer (DeNovix). First-strand cDNA was produced in a 20 µL reaction volume using the Superscript IV kit (Invitrogen).

The cDNA was diluted with water in a 1:20 ratio and 2 µL of this solution was used for qRT-PCR in a 7 µL reaction volume using a EvaGreen Mastermix (Metabion, UNG+/ROX+ 2x conc.) and primers shown in Supplementary Table 2. To quantify the expression of the different genes, the qPCR reaction was carried out using a CFX384 Touch^TM^ RT-PCR detection system (Bio-Rad). Thermal cycler conditions were: 95°C 2 min, 40 cycles of 95°C 30s, 55°C 30s and 72°C 20 s, followed by dissociation curve analysis. For the calculation of the expression levels, we followed the ΔΔCt method (Czechowski et al., 2004). For each genotype three biological replicates were analyzed. Each sample was represented by 3 technical replicates.

### Statistical analysis

Statistical analyses were performed in R-studio, using one-way Analysis of Variance (ANOVA), followed by Tukey HSD post hoc test.

### Accession numbers

Sequence data for the genes mentioned in this article can be found in The Arabidopsis Information Resource (TAIR; https://www.arabidopsis.org) under the following accession numbers: *MAX3*, AT2G44990; *MAX4*, AT4G32810; *MAX1*, AT2G26170; *D14*, AT3G03990; *KAI2*, AT4G37470; *MAX2*, AT2G42620; *SMAX1*, AT5G57710; *SMXL2* AT4G30350; *SMXL6*, AT1G07200; *SMXL7*, AT2G29970; *SMXL8*, AT2G40130; *PIN1*, AT1G73590; *PIN2*, AT5G57090; *PIN3*, AT1G70940; *PIN4*, AT2G01420; *PIN7*, AT1G23080; *ARF5*, AT1G19850; *ARF7*, AT5G20730; *ARF8*, AT5G37020; *ARF19*, AT1G19220; *RSL2*, AT4G33880; *RSL4*, AT1G27740; *EXP7*, AT1G12560; *COW1*, AT4G34580; *DLK2*, AT3G24420, *TAA1*, AT1G70560; *DAO1*, AT1G14130, *DAO2*, AT1G14120; *YUC3*, AT1G04510; *YUC9*, AT1G04180; *TIR1*, AT3G62980; *AUX1*, AT2G38120; *EF1α*, AT5G60390.

## AUTHOR CONTRIBUTIONS

JAVA, TB and CG conceived the study, designed experiments, analyzed and interpreted data. JAVA performed all root hair, skewing, waving, cell length and root diameter analyses (except for Supplementary Figure 3B, 3B, 6C and Figure 6C, 6D, 7D and 7D), gene expression analysis, prepared figures, performed statistics and wrote the first draft of the abstract, figure legends and materials and methods. MHJ performed all PRL, LRD and skewing (Supplementary Figure 6C) analysis in [L] and quantified *DR5v2:GFP* expression. SC acquired the data for Figure 6C, 6D, 7D and 7D, and AK for Supplementary Figure 3B and 3D. KL quantified IAA levels (Figure 9A). TB performed PRL, LRD and skewing (data not shown) analysis in [C] as well as PIN:GFP quantification. KL, TB and CG acquired funding. TB and CG were PIs of the study and wrote the manuscript.

## ACKNOWLEDGEMENTS

We thank David Nelson (UC Riverside, USA) and Mark Waters (University of Western Australia) for providing mutant seeds. The Gutjahr group is grateful to Jürgen Soll (LMU Munich, Germany) for generously providing space in his *Arabidopsis* growth chamber, to Regina Hüttl for excellent technical assistance and Patrick Wiesbeck for help with root hair quantification. KL thanks Roger Granbom for excellent technical assistance, and the Swedish Research Council (VR) and the Swedish Governmental Agency for Innovation Systems (VINNOVA) for funding. This work was supported by the BBSRC grant BB/R00398X/1 to TB and by the Emmy Noether program of the Deutsche Forschungsgemeinschaft (DFG) to CG (GU1423/1-1).

## Supplemental Data

The following materials are available in the online version of this article.

**Supplemental Figure 1.** Strigolactone signalling regulates primary root length and lateral root

**Supplemental Figure 2**. KL signalling regulates lateral root and root hair development.

**Supplemental Figure 3.** KL perception mutants respond to tilted agar surface.

**Supplemental Figure 4.** KL perception mutants exhibit decreased epidermal cell lengths and decreased root diameter in Ler background.

**Supplemental Figure 5.** Root hair development is not regulated by the SMXL678.

**Supplemental Figure 6.** KAI2 regulation of root skewing can be genetically separated from root diameter.

**Supplemental Figure 7.** GR24 stereoisomers regulate root hair development through D14 and KAI2.

**Supplemental Figure 8.** Auxin biosynthesis and transport gene expression in KL perception mutants.

**Supplemental Table 1.** Summary of effects of *SMXL* mutations on *max2* root phenotypes

**Supplemental Table 2.** List of primers used for qPCR analysis

