## supplemental material for "KAI2 regulates root and root hair development by modulating auxin distribution" (from another version of this preprint: version 1, 2019-02-03)

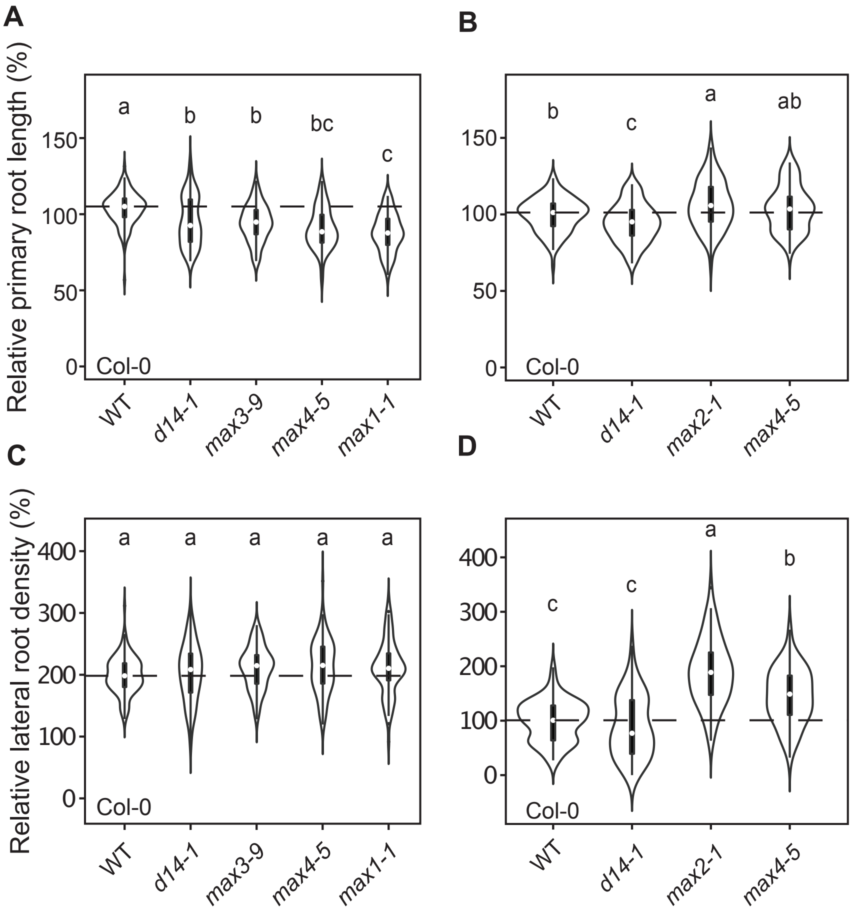


**Supplementary Figure 1. Primary root and lateral root development in SL perception and biosynthesis mutants.**

**(A, B)** Relative primary root length and **(C** and **D)** relative lateral root density of strigolactone perception and biosynthesis mutants. Primary root length and lateral root density are presented relative to WT (in %). The outline of the violin plots represents the probability of the kernel density. Black boxes represent interquartile ranges (IQR), with the white dot representing the median; whiskers extend to the highest and lowest data point but no more than ±1.5 times the IQR from the box; outliers are plotted individually. Different letters indicate different statistical groups (ANOVA, posthoc Tukey, **(A)** $F_{5,503} = 19.75$; $p \leq 0.001$, **(B)** $F_{4,387} = 10.71$; $p \leq 0.001$, **(C)** $F_{5,503} = 1.985$; $p < 0.05$, **(D)** $F_{4,257} = 39.66$; p0.05).

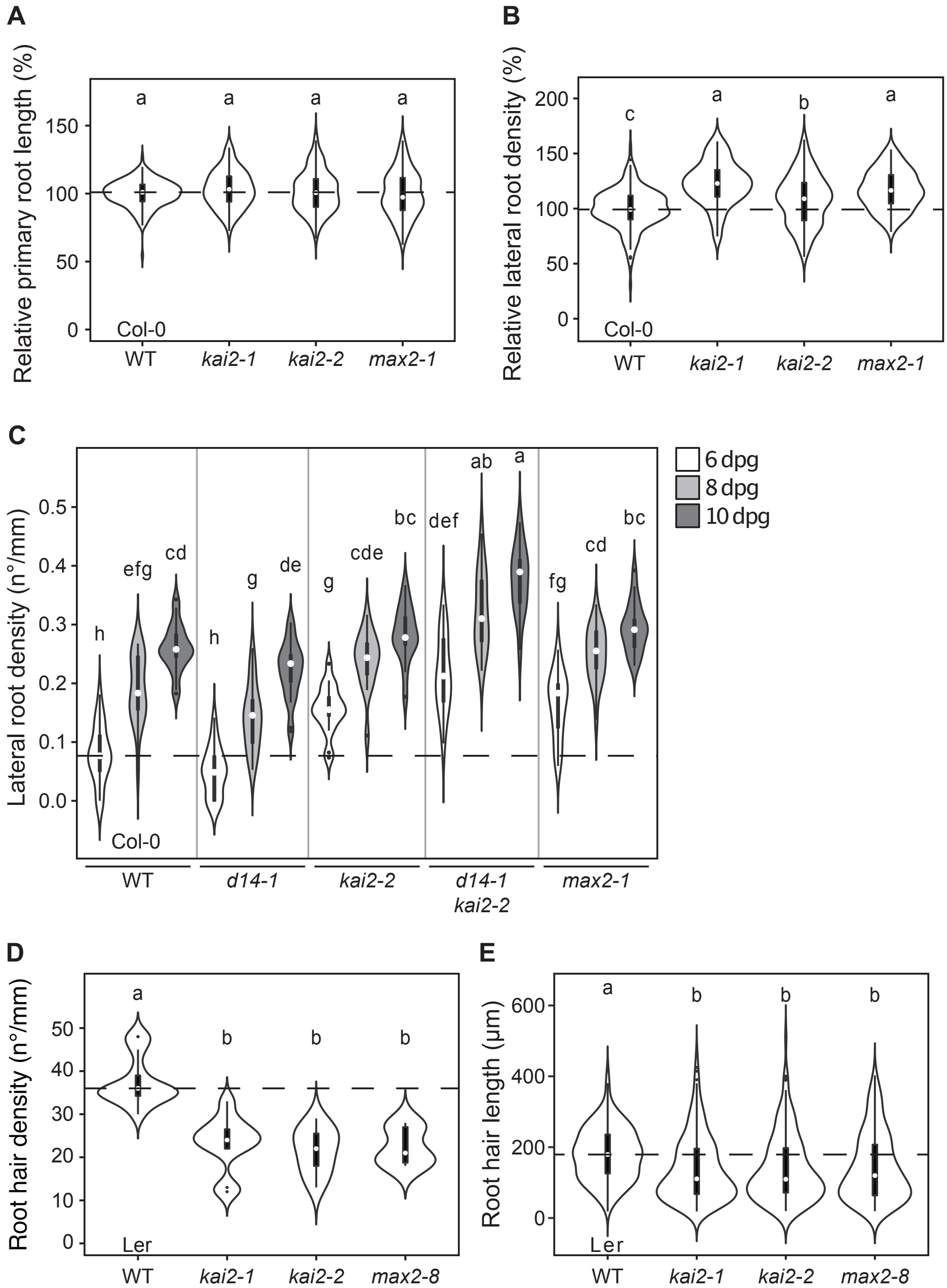


**Supplementary Figure 2. KL signaling regulates lateral root and root hair development**
**(A)** Relative primary root length and **(B)** relative lateral root density of karrikin perception mutants. Primary root length and lateral root density are presented relative to WT (in %). **(C)** Lateral root density at 6, 8 or 10 days post germination (dpg), **(D)** root hair density and **(E)** root hair length of the indicated genotypes. The outline of the violin plots represents the probability of the kernel density. Black boxes represent interquartile ranges (IQR), with the white dot representing the median; whiskers extend to the highest and lowest data point but no more than ±1.5 times the IQR from the box; outliers are plotted individually. Different letters indicate different statistical groups (ANOVA, posthoc Tukey, **(A)** $F_{3,520} = 1.381$; $p<0.05$, **(B)** $F_{3,520} = 29.86$; $p \leq 0.001$, **(C)** $F_{4,295} = 33.79$; $p \leq 0.001$, **(D)** $F_{3,754} = 10.83$; $p \leq 0.001$, **(E)** $F_{3,40} = 24.27$; $p \leq 0.001$).

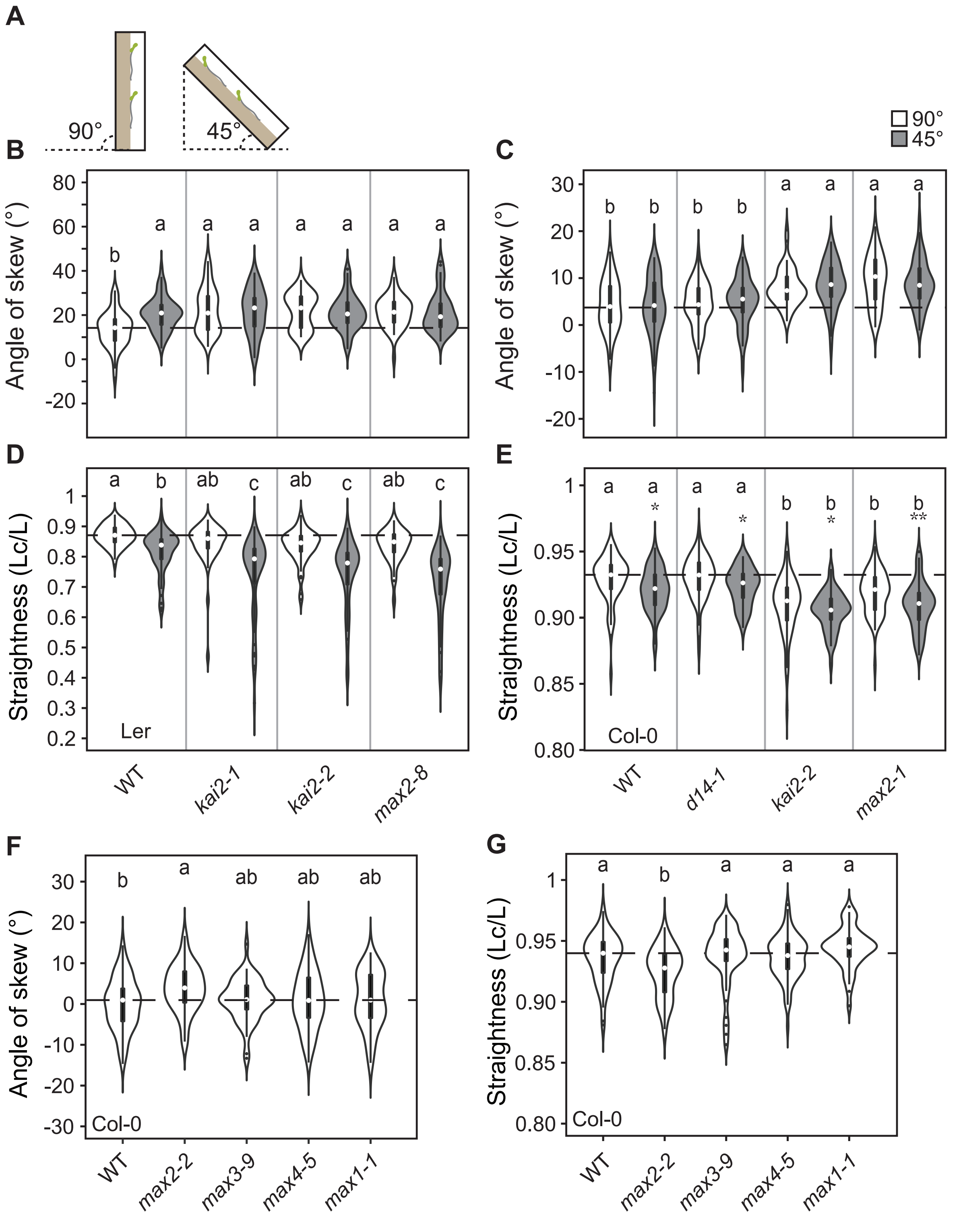


**Supplementary Figure 3. KL perception mutants respond to tilted agar surface.**
**(A)** Diagram showing agar plates tilted at 90° or 45° from the vertical. Root skewing **(B, C, F)** and root straightness **(D, E, G)** of the indicated genotypes. Plants were grown at 90° or 45° from the vertical **(A, B, C, D, E)** or 90° from the vertical **(F, G)**. The outline of the violin plots represents the probability of the kernel density. Black boxes represent interquartile ranges (IQR), with the white dot representing the median; whiskers extend to the highest and lowest data point but no more than ±1.5 times the IQR from the box; outliers are plotted individually. Different letters indicate different statistical groups (ANOVA, posthoc Tukey, **(B)** $F_{7,383} = 5.788$; $p \leq 0.001$, **(C)** $F_{7,472} = 12.54$; $p \leq 0.001$, **(D)** $F_{7,430} = 25.89$; $p \leq 0.001$, **(E)** $F_{7,497} = 18.36$; $p \leq 0.001$, **(F)** $F_{5,333} = 5.057$; $p \leq 0.001$, **(G)** $F_{4,290} = 7.168$; $p \leq 0.001$).

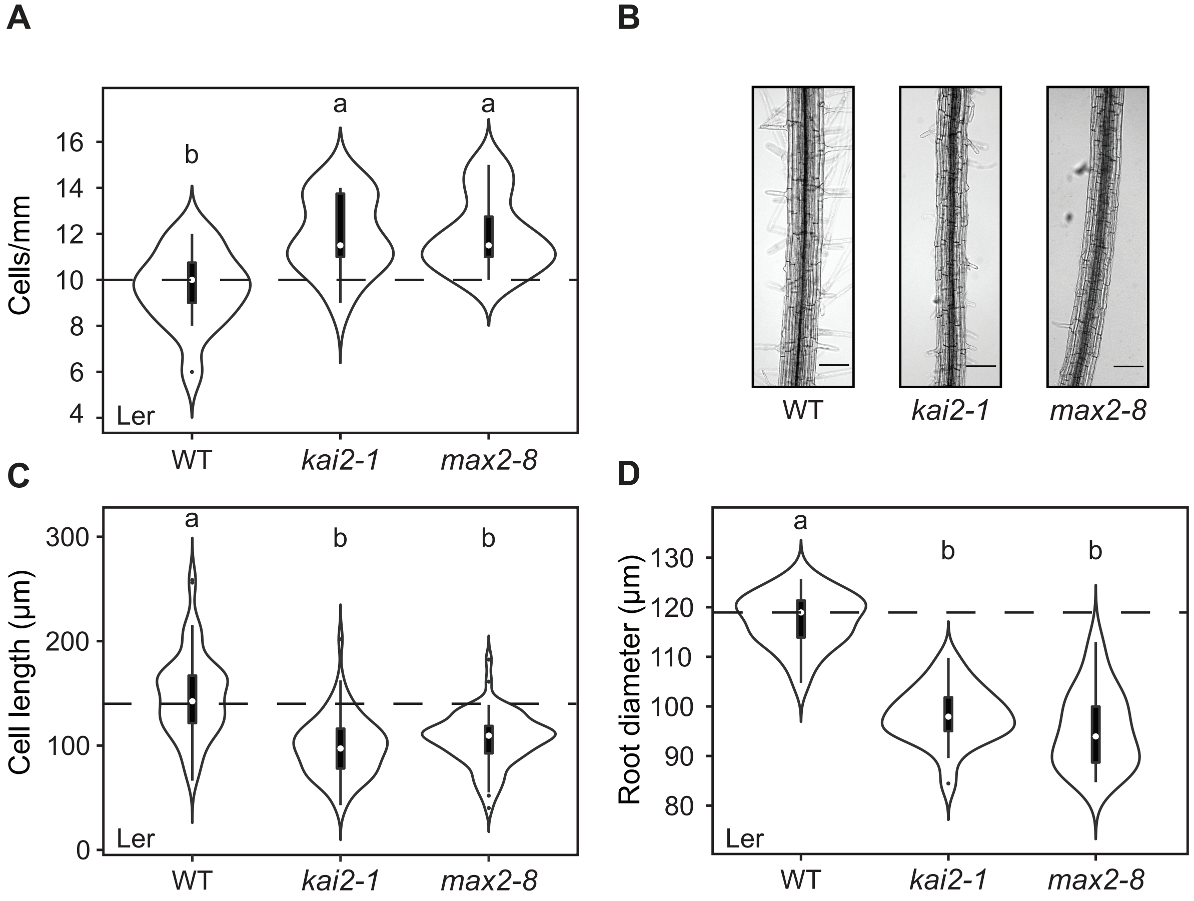


**Supplementary figure 4. KL perception mutants exhibit decreased epidermal cell lengths and increased root diameter in Ler background.**

**(A)** Number of root epidermal cells per mm of the indicated genotypes. **(B)** Images of representative roots between 2 and 3 mm from the root tip from 5-days-old seedlings. Scale bars, 0.1 mm. **(C)** Root cell length and **(D)** root diameter of the indicated genotypes. The outline of the violin plots represents the probability of the kernel density. Black boxes represent interquartile ranges (IQR), with the white dot representing the median; whiskers extend to the highest and lowest data point but no more than ±1.5 times the IQR from the box; outliers are plotted individually. Different letters indicate different statistical groups (ANOVA, posthoc Tukey, **(A)** $F_{2,43} = 9.58$; $p \leq 0.001$, **(C)** $F_{2,191} = 43.1$; $p \leq 0.001$, **(D)** $F_{2,64} = 77.45$; $p \leq 0.001$).

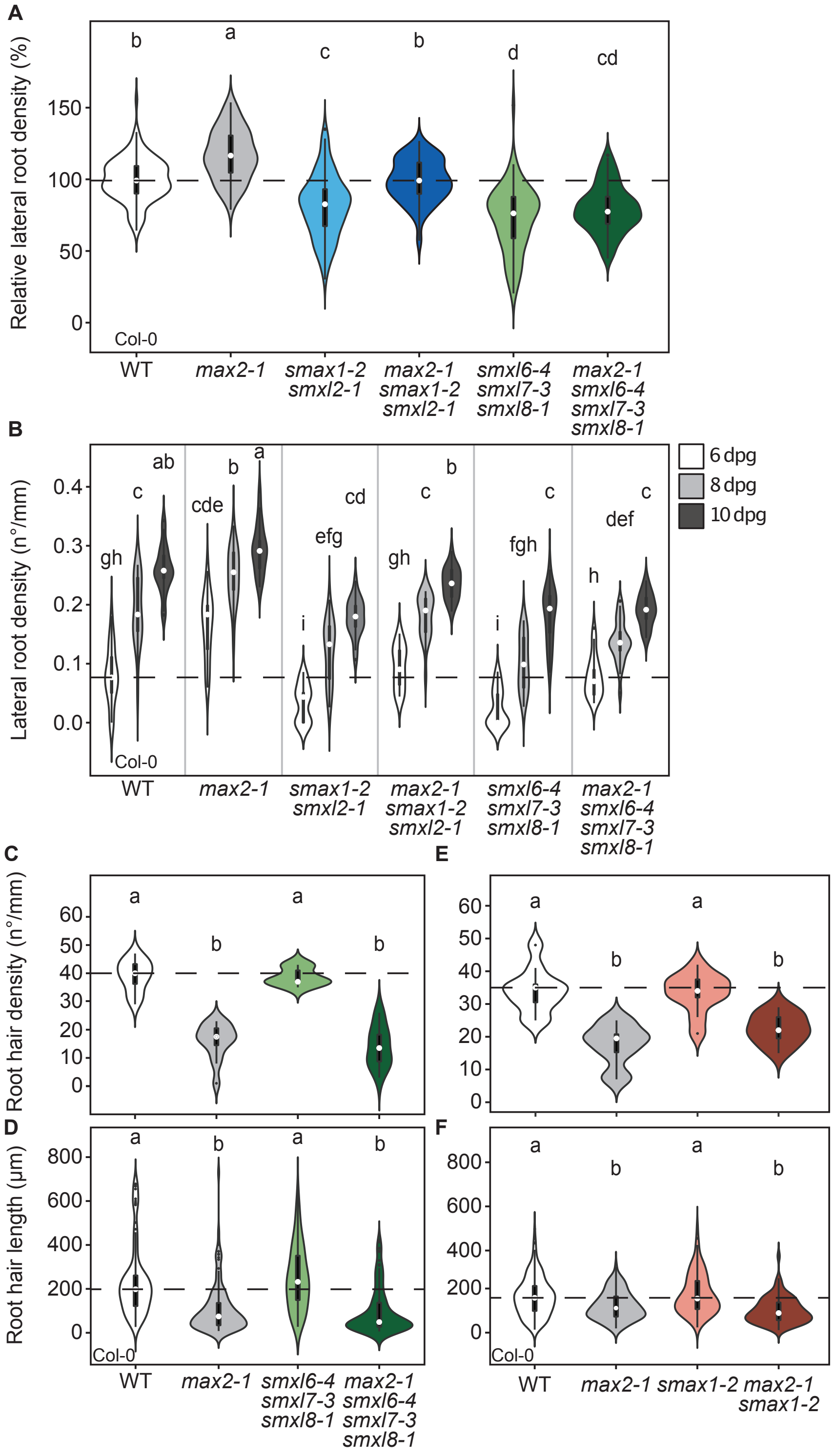


**Supplementary Figure 5. Lateral root and root hair development are not regulated by the SL signalling repressors SMXL6,7,8.**

**(A)** Relative lateral root density of the indicated genotypes. Lateral root density is presente relative to the WT (in %). **(B)** lateral root density at 6, 8 or 10 days post germination (dpg), **(C, E)** root hair density and **(D, F)** root hair length of the indicated genotypes. The outline of the violin plots represents the probability of the kernel density. Black boxes represent interquartile ranges (IQR), with the white dot representing the median; whiskers extend to the highest and lowest data point but no more than ±1.5 times the IQR from the box; outliers are plotted individually. Different letters indicate different statistical groups (ANOVA, posthoc Tukey, **(A)** $F_{5,580} = 79.32$; $p \leq 0.001$, **(B)** $F_{17,385} = 79.17$; $p \leq 0.001$, **(C)** $F_{3,44} = 67.45$; $p \leq 0.001$, **(D)** $F_{3,410} = 38.66$; $p \leq 0.001$, **(E)** $F_{3,39} = 20.33$; $p \leq 0.001$, **(F)** $F_{3,615} = 30.02$; $p \leq 0.001$).

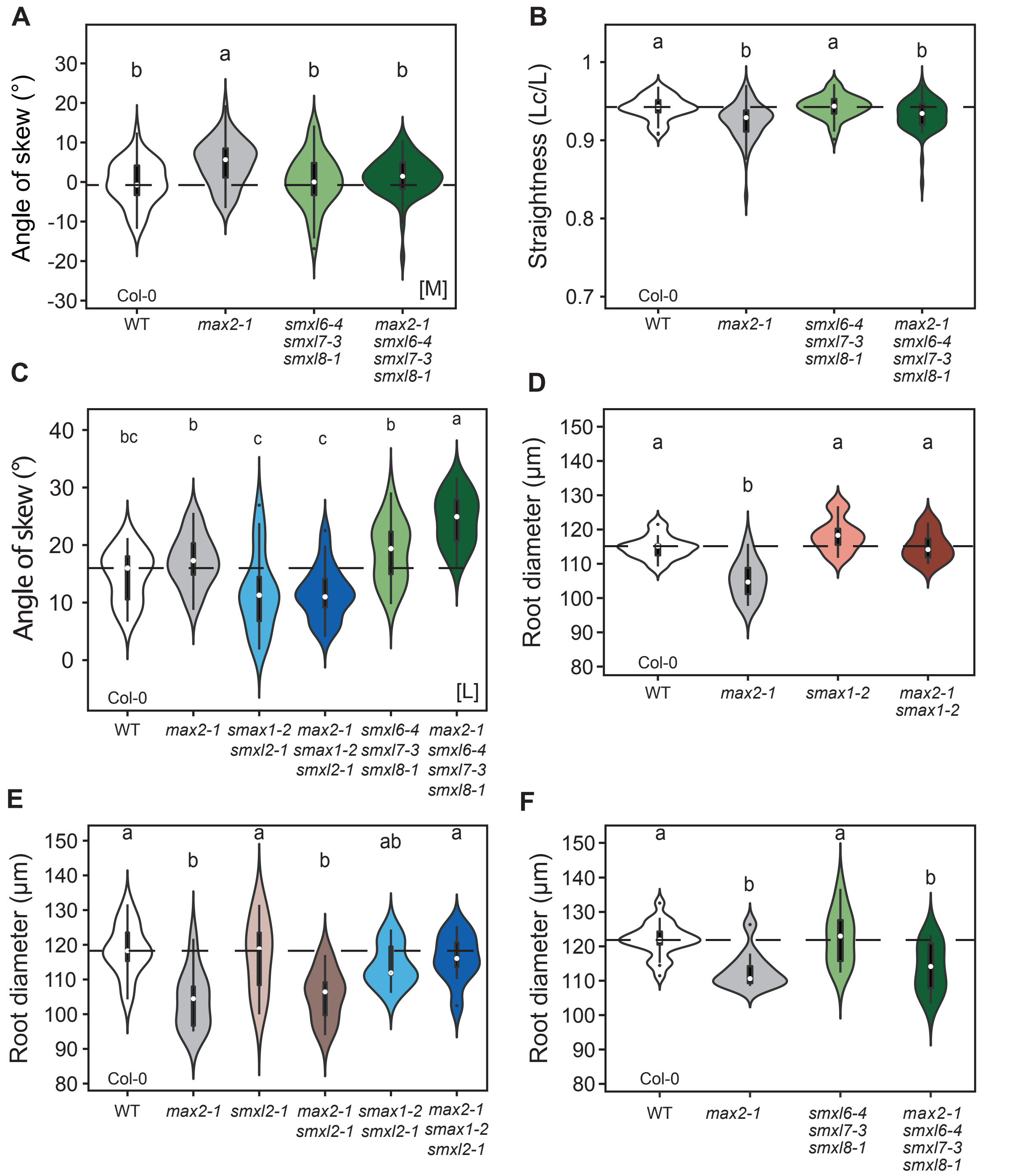


**Supplementary Figure 6. KAI2 regulation of root skewing can be genetically separated from root diameter.**
**(A, C)** Root skewing, **(B)** root straightness and **(D, E, F)** root diameter of the indicated genotypes. The outline of the violin plots represents the probability of the kernel density. Black boxes represent interquartile ranges (IQR), with the white dot representing the median; whiskers extend to the highest and lowest data point but no more than ±1.5 times the IQR from the box; outliers are plotted individually. Different letters indicate different statistical groups (ANOVA, posthoc Tukey, **(A)** $F_{3,209} = 8.784$; $p \leq 0.001$, **(B)** $F_{3,209} = 10.22$; $p \leq 0.001$, **(C)** $F_{5,127} = 21.07$; $p \leq 0.001$, **(D)** $F_{3,38} = 15.04$; $p \leq 0.001$, **(E)** $F_{3,38} = 15.04$; $p \leq 0.001$, **(F)** $F_{3,47} = 8.221$; $p \leq 0.001$). [M] = Munich, [L] = Leeds.

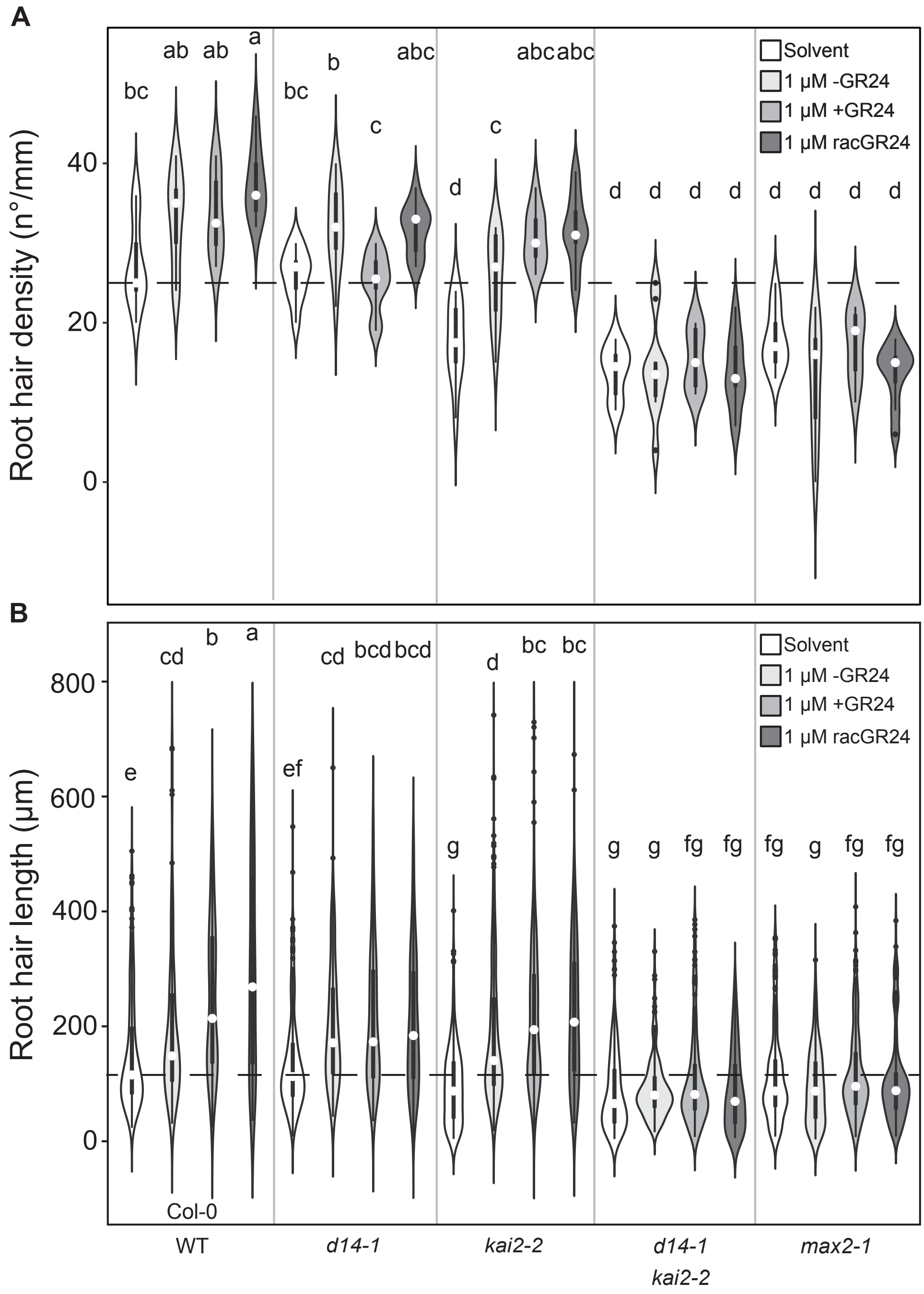


**Supplementary Figure 7. GR24 stereoisomers regulate root hair development through D14 and KAI2.**

**(A)** Root hair density and **(B)** root hair length of the indicated genotypes treated with solvent (acetone), 1 µM µM –GR24, 1 µM +GR24 or 1 µM *rac*-GR24. The outline of the violin plots represents the probability of the kernel density. Black boxes represent interquartile ranges (IQR), with the white dot representing the median; whiskers extend to the highest and lowest data point but no more than ±1.5 times the IQR from the box; outliers are plotted individually. Different letters indicate different statistical groups (ANOVA, posthoc Tukey, **(A)** $F_{19, 3740} = 1.983$; $p \leq 0.01$, **(B)** $F_{19,3740} = 57.83$; $p \leq 0.001$).

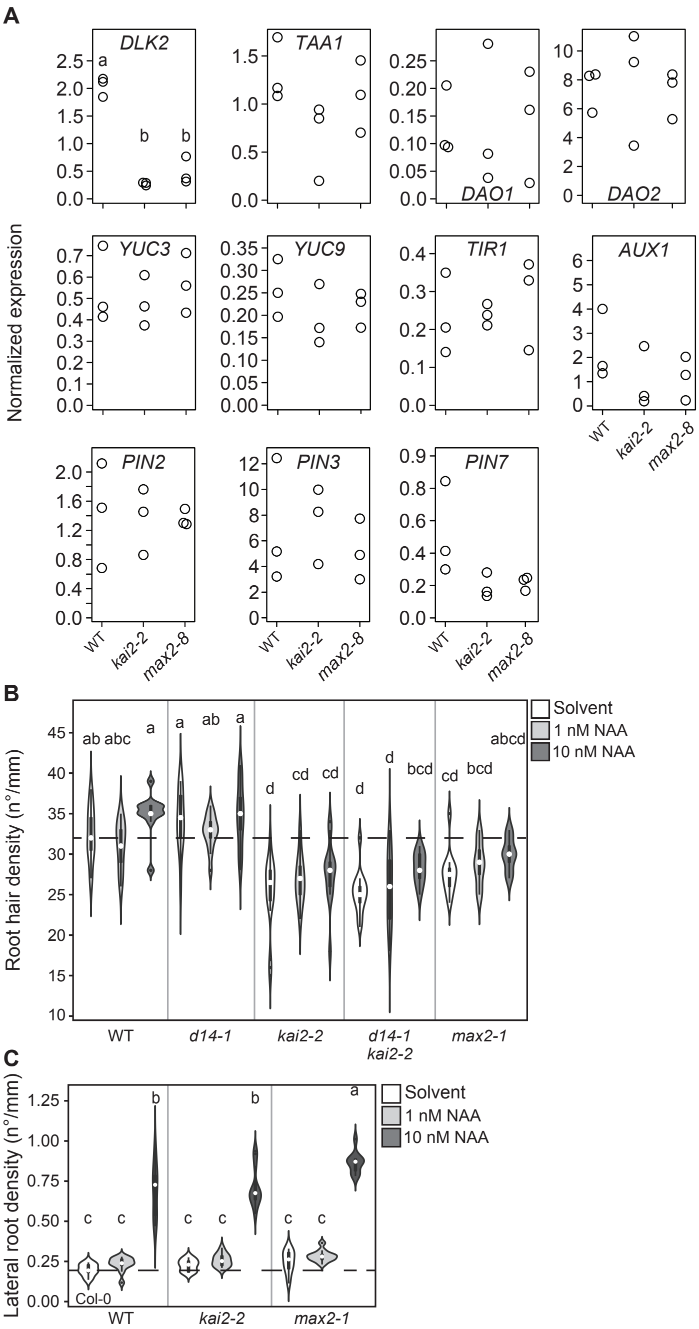


**Supplementary Figure 8. Auxin biosynthesis and transport gene expression**

**(A)** Transcript accumulation of *DLK2*, *TAA1*, *DAO1*, *DAO2*, *YUC3*, *YUC9*, *TIR1, AUX1, PIN2*, *PIN3* and *PIN7* in roots of the indicated genotypes. Expression levels of 3 biological replicates are normalized against those of *EF1α*. **(B)** Root hair density and **(C)** lateral root density of the indicated genotypes, treated with solvent (96% Ethanol), 1 nM NAA or 10 nM NAA. The outline of the violin plots represents the probability of the kernel density. Black boxes represent interquartile ranges (IQR), with the white dot representing the median; whiskers extend to the highest and lowest data point but no more than ±1.5 times the IQR from the box; outliers are plotted individually. Different letters indicate different statistical groups (ANOVA, posthoc Tukey, **(A)** $F_{2,6} = 90.81$; $p \le 0.001$ (*DLK2*), $F_{2,6} = 2.365$; $p < 0.05$ (*TAA1*), $F_{2,6} = 0.005$; $p \le 0.05$ (*DAO1*), $F_{2,6} = 0.061$; $p \le 0.05$ (*DAO2*), $F_{2,6} = 0.061$; $p < 0.05$ (*YUC3*), $F_{2,6} = 0.906$; $p < 0.05$ (*YUC9*), $F_{2,6} = 1.018$; $p < 0.05$ (*TIR1*), $F_{2,6} = 1.281$; $p < 0.05$ (*AUX1*), $F_{2,6} = 0.906$; $p < 0.05$ (*PIN2*), $F_{2,6} = 0.332$; $p < 0.05$ (*PIN3*), $F_{2,6} = 3.297$; $p < 0.05$ (*PIN7*), **(B)** $F_{5,63} = 31.25$; $p \le 0.001$, **(C)** $F_{5,67} = 24.69$; $p \le 0.001$).

**Supplemental Table S1.** Summary of effects of *SMXL* mutations on *max2* root phenotypes.

| | *max2* | | | |
|---|---|---|---|---|
| | *smax1* | *smxl2* | *smax1 smxl2* | *smxl6,7,8* |
| Lateral root density | | | + | + |
| Root hair density | - | - | + | - |
| Root hair length | - | - | + | - |
| Root skewing | + | + | + | +/- |
| Root straightness | - | - | + | - |
| Root diameter | + | - | + | - |

\+ Suppression of *max2* phenotypes

\- No suppression of *max2* phenotypes

+/- Opposite effects in Munich (+) and Leeds (-)

**Supplemental Table 2.** List of primers used for qPCR analysis.

| Gene | Direction | Sequence | References |
|---|---|---|---|
| *ARF5* | Forward | TACGCCACTTAGAACCAGGCCATC | |
| | Reverse | GTTCCCATGTACCTTCGTTTCCCG | |
| *ARF7* | Forward | GCTCATATGCATGCTCCACA | Singh et al., 2012 |
| | Reverse | CAGCAGGAGCAGCCCACCT | |
| *ARF8* | Forward | ACATGGAGGGTTTTCTGTTCC | |
| | Reverse | TGGCACTGACAAAGACACTCCATC | |
| *ARF19* | Forward | TCTTGCAGCTATCCCAACCA | |
| | Reverse | CGATGGCCTCGAATGATAATGTAA | |
| *RSL2* | Forward | TCCCCAATGGAACAAAGGTC | Yi et al., 2010 |
| | Reverse | TCTCGGTGAGCTGAGACCAA | |
| *RSL4* | Forward | GTGCCAAACGGGACAAAAGT | Yi et al., 2010 |
| | Reverse | TTGTGATGGAACCCCATGTC | |
| *EXP7* | Forward | AACCATGGGTGGTGCATG | Zhang et al., 2016 |
| | Reverse | CCGCATCCGTAACCATCA | |
| *COW1* | Forward | CCACATGATGCTTCGATTTTTGAG | Zhang et al., 2016 |
| | Reverse | TAGCCTTGAGGGTAGTGC | |
| *DLK2* | Forward | GCTGCTTCTCCAAGGTATATAA | Waters et al., 2012 |
| | Reverse | GAAATCAACCGCCCAAGCT | |
| *TAA1* | Forward | ATCTTACCCTGCGTTTGCGT | |
| | Reverse | AGCATGCTGACTCGGACATGC | |
| *DAO1* | Forward | CTGCAGATCAAAGGGAGATT | |
| | Reverse | TCTCAACCAGCCCGTAACTC | |
| *DAO2* | Forward | TGGTGACATGGCTACGATATG | |
| | Reverse | CTTCAAGATCTCTATCCACTGG | |
| *YUC3* | Forward | TCGTAGCGCTGTTCATGTTT | |
| | Reverse | GCGAGCCAAACGGGCATATACTTC | Liu et al., 2017 |
| *YUC9* | Forward | AGTCCGGCGAGAAATTCAGAGG | Liu et al., 2017 |
| | Reverse | AACATGAACCGAGCTTCTAACGAC | |
| *TIR1* | Forward | GCCACTTGCAGGAATCTGAA | |
| | Reverse | TGAGAGACTTGAGATTGGGACA | |
| *AUX1* | Forward | GCTGTCGGTGCTCTTCTTG | |
| | Reverse | CTTCTCCGCCGCATTCTGA | |
| *PIN2* | Forward | TCACGACAACCTCGCTACTAAAGC | Niu et al., 2015 |
| | Reverse | GTCTTGGTCCATTTCCACATGCC | |
| *PIN3* | Forward | GAGCACCTGACAACGATCAAGG | Niu et al., 2015 |
| | Reverse | GATGAGCTACAGCTTTGGTC | |
| *PIN7* | Forward | GGAGCCAATGAACAAGTCGG | D´Alessandro et al., 2015 |
| | Reverse | TCATCGGACCAGCTTTGTTT | |
| *EF1α* | Forward | GGTGGTGGCATCCATCTTGTTACA | Yi et al., 2010 |
| | Reverse | TGAGCACGCTCTTCTTGCTTTCA | |
